# Adsorbing DNA to mica by cations: Influence of the valency and ion type

**DOI:** 10.1101/2023.06.30.547224

**Authors:** Mohd Ibrahim, Christiane Wenzel, Max Lallemang, Bizan N. Balzer, Nadine Schwierz

## Abstract

Ion-mediated attraction between DNA and mica plays a crucial role in biotechnological applications and molecular imaging. Here, we combine molecular dynamics simulations and single-molecule atomic force microscopy experiments to characterize the detachment forces of single-stranded DNA at mica surfaces mediated by the metal cations Li^+^, Na^+^, K^+^, Cs^+^, Mg^2+^ and Ca^2+^. Ion specific adsorption at the mica/water interface compensates (Li^+^, Na^+^) or overcompensates (K^+^, Cs^+^, Mg^2+^ and Ca^2+^) the bare negative surface charge of mica. In addition, direct and water-mediated contacts are formed between the ions, the phosphate oxygens of DNA and mica. The different contact types give rise to low and high force pathways and a broad distribution of detachment forces. Weakly hydrated ions, such as Cs^+^ and water-mediated contacts lead to low detachment forces and a high mobility of the DNA on the surface. Direct ion-DNA or ion-surface contacts lead to significantly higher forces. The comprehensive view gained from our combined approach allows us to highlight the most promising cations for imaging in physiological conditions: K^+^ to overcompensate the negative mica charge and induce long-ranged attractions. Mg^2+^ and Ca^2+^ to from a few specific and long-lived contacts to bind DNA with high affinity.

## Introduction

The adsorption of nucleic acids to solid substrates plays a crucial role for molecular imaging and a variety of biotechnological applications such as biosensors and micro-arrays.^1–5^ Atomic force microscopy (AFM) has been applied in countless studies to image and characterize nucleic acids, DNA-protein complexes or DNA nano-structures.^6–14^ Muscovite mica, a highly negatively charged and atomically smooth surface, is one of the most popular substrates for such biomolecular imaging. ^5,15–21^ However, the negative charge of mica and DNA hinders adsorption and metal cations are essential for adsorption and immobilization. The influence of different cations on DNA-mica interactions has hence been studied intensively using AFM imaging techniques.^5,7,15–21^ Divalent transition metals have proven to be particularly useful to immobilize the DNA.^15–17^ Transition metals (Ni^2+^, Zn^2+^, Co^2+^) can strongly bind DNA, and the binding affinity increases with charge density, whereas alkaline earth metals such as Mg^2+^ and Ca^2+^ are less effective.^15,16^ While transition metals have the benefit of high signal-to-noise images, they have two major drawbacks when considering the physiologically relevant properties and interactions of biomolecules: Due to the high binding affinity, DNA is kinetically trapped in a collapsed conformation on the surface. ^22^ In the presence of divalent metals, the conformation of DNA or DNA-protein complex likely differs from the one in physiological conditions.^23^ Recently, Heenan and Perkins ^5^ successfully imaged DNA on mica in solutions containing biochemically relevant conditions of Mg^2+^ and K^+^ to preserve native bio-molecular interactions. However, the molecular interactions that lead to preferential interactions have not been resolved and it remains unknown how to choose the ionic condition and the type of ion in solution to tune the binding strength.

It has been known for a long time that like-charged objects, such as DNA and mica in the present case, can attract each other in the presence of multivalent cations. Due to its importance, the theoretical modelling of the adsorption of charged polymers at interfaces has steadily progressed.^24^ However, traditional approaches based on Debye-Hückel or Poisson-Boltzmann theory predict purely repulsive forces between similarly charged objects. The failure is typically attributed to the lack of ion correlations in the theory. Ion correlations originate from three different sources:^25^ (i) ion-density fluctuations around the mean ion distribution, (ii) non-electrostatic ion-ion interactions and (iii) direct and water-mediated interactions between the ions and the surfaces. Including the condensation of the cations at highly charged polymers via counterion condensation theory predicts an attractive regime for mono- and divalent ions.^26^ Similarly, the Oosawa model, which includes ion-density fluctuations, predicts an attractive regime for divalent ions.^27^ Numerous similar approaches which attempt to treat correlations on a more fundamental level exist in the literature.^25,28–31^ However, it is challenging to apply such simple models to highly charged biological systems and to quantitatively predict the interactions between ions, mica, and DNA. Firstly, the Coulomb interactions are long-ranged, couple many charged objects and yield a manybody problem. Secondly, DNA molecules are flexible and their exact conformation depends on the electrostatic environment.^32^ Finally, the adsorption of ions at the mica/water interface and at the DNA molecule depends on the type of cation.^33,34^ Even though such ion-specific effects can be included into Poisson-Boltzmann (PB) theory,^31,35^ there is no framework that allows us to quantitatively predict the forces between DNA and mica due to the entangled contributions of ion specific adsorption of DNA and mica and hydration effects.

All-atom molecular dynamics simulations in explicit water are ideal to resolve all individual contributions. Moreover, combined with single-molecule experiments they quantify the adhesion forces of biomolecules and provide insights into the mechanism of ion-mediated adsorption. Similarly, force spectroscopy experiments are versatile methods to measure the molecular interaction forces in the range of piconewtons^36,37^ and have been successfully applied for biomolecules at lipid bilayers, gold and graphite surfaces.^36,38–43^ Steered molecular dynamics simulations closely mimic this experimental setup. The direct comparison of the desorption forces, albeit challenging due to the different timescales accessible,^44^ allow us to verify the simulations or to uncover shortcoming of the atomistic force fields.^32,38,39,45–47^

In this current work, we combine all-atom molecular dynamics simulations and AFM experiments to characterize the detachment forces of single-stranded DNA from muscovite mica in the presence of the mono- and divalent metal ions Li^+^, Na^+^, K^+^, Cs^+^, Mg^2+^ and Ca^2+^. Initially, we characterize the ion-specific interfacial structure at the mica/electrolyte interface. Subsequently, we directly compare detachment forces as a function of the loading rate and shape of force-extension curves from simulations and experiments. Finally, we provide the distribution of detachment forces for all ions and resolve the molecular interactions that contribute to high and low forces. The distribution of DNA-mica forces for different metal ions provides a baseline of how to choose ionic conditions to tune the binding strength of DNA.

## Materials and Methods

### Simulations

#### Setup

Muscovite mica (KAl_3_Si_3_O_10_(OH)_2_) was simulated with dimensions 8.264 *×* 7.172 *×* 4.0 nm^3^. The 001 plane was set orthogonal to the z-axis of the simulation box. The topmost layer of K^+^ ions was removed leading to negatively charged surfaces (*σ*_bare_ = *−*2.16 e/nm^2^). A single-stranded DNA (poly-DT_10_) was placed at a distance of about 2 nm from the mica surface. The systems were solvated in TIP3P water.^48^The number of water molecules in the simulation box were between 21900-26800. This corresponds to a box size in the z-dimension between 14.4 −16.7 nm. Ions were added to obtain a bulk concentration of 150 mM. In particular, 327 Li ^+^, 325 Na^+^, 360 K^+^, 409 Cs^+^, 214 Mg^2+^ and 212 Ca^2+^ cations, and the respective number of Cl^*−*^ ions were added to yield neutral systems and to obtain the correct bulk concentration (Figure S1). Note that the different number of cations reflects the ion-specific binding affinities of the cations to mica.^33^ The resulting effective surface charge (*σ*_eff_) is listed in Table 1 (see supporting information section 1.2, Figure S2). The adjustment procedure to obtain a bulk concentration of 150 mM is described in the supporting information(section 1.1, Figure S1).

The parmBSC1 forcefield^49^ was used for DNA and CLAYFF forcefield^50^ was used to describe mica. We chose the Mamatkulov-Schwierz force field parameters^51^ for the monovalent ions since they were optimized based on experimental hydration free energies and activity coefficients. In addition, these force fields were shown to correctly reproduce oscillatory hydration forces at mica surfaces for monovalent cations.^33^ For divalent Mg^2+^ and Ca^2+^ recently developed force field parameters were used^52,53^ (nanoMg parameter set). These parameters reproduce experimental hydration free energies, activity coefficients, water exchange rates and the binding affinities to the phosphate oxygens on the backbone of nucleic acids. Since the DNA force field and the optimized ion parameters work in combination with TIP3P, this water model was used in our current work, even though other models reproduce the physical properties of bulk water better compared to TIP3P.^54^

We first performed an energy minimization. Subsequently, the systems were equilibrated for 50 ns in the NAP_*z*_T ensemble using semi-isotropic pressure coupling via the Berendsen barostat and thermostat.^55^ We applied position restraints on the heavy atoms of the DNA such that ions can adsorb and form an electric double layer prior to DNA adsorption. Pressure and temperature were fixed at 1 bar and 300 K respectively using a time constant of 1 ps. Subsequently, the restraints on the DNA were removed and the system was relaxed for 1 ns. For Mg^2+^, we used the nanoMg parameter set.^53^ With this parameter set, we could not observe any inner-sphere binding of Mg^2+^ on the nanosecond timescale. However, due to the presence of high-force pathways in the experiments, we conclude that inner-sphere conformations must exist. We therefore performed additional high temperature simulations at 600 K for 100 ns. At 600 K, Mg^2+^ sheds off its hydration shell and forms inner-sphere contacts with the DNA. Subsequently, the temperature was reduced to 300 K. After equilibration, the pulling was performed as described below.

#### Detachment forces from simulations

The simulation routine consisted of the following steps. (i) Initially the DNA was forced to adsorb at the surface by applying a constant external force *F*_ext_ to all heavy atoms of the DNA. This forced adsorption mimics the AFM setup, in which a force of approx. 500 pN is used to bring the DNA to the mica/water interface. To provide additional information on the adsorption behavior, we used different values for the external force in the range of 0 *−* 470 pN (see supporting information, section 1.3, Figure S3). The simulations were performed for 50 ns and the work of adhesion was calculated similarly as in previous work. ^39^ In the absence of an external force, the DNA spontaneously adsorbed at the mica/water interface (K^+^, Cs^+^) or remained in bulk water (Li^+^, Na^+^). This indicates that in the presence of Li^+^ and Na^+^ the strongly bound hydration layer prevents spontaneous adsorption similar to hydrophilic surfaces.^38^ Finally, the highest force value *F*_ext_ = 470 pN was selected for all systems.

(ii) Subsequently, the external force was removed and the system was equilibrated for 230-300 ns. After about 100 ns, the number of contacts and the distance of the DNA from the mica surface remained constant (Figure S4). The first 100 ns were therefore discarded for equilibration. For low pulling velocities, we selected six configurations every 20 ns and 130 conformations every 1 ns for the highest pulling velocity as starting points for the dynamic pulling simulations. (iii) Finally, dynamic pulling simulations were performed. The pulling simulations closely mimic the AFM experiments and the DNA was pulled away perpendicular to the mica surface with constant velocity using the Gromacs pull code.^56^ To mimic the constant velocity experiments, a harmonic potential was applied to the center of mass of the 5^*′*^-end residue of DNA with respect to the center of mass of mica surface atoms and moved away from the surface with constant velocity.

We used 3 different pulling velocities: 10 m/s, 1.0 m/s, and 0.1 m/s. For *v* = 10 m/s, 100-150 pulling simulations were performed with different starting configurations and 6 otherwise. The spring constant was fixed at *k*_*s*_ = 166 pN/nm similar to previous work.^38,47^ Note that this value is a factor of 5 to 10 higher than in the AFM experiments. The resulting force-extension curves for *v* = 0.1 m/s for all ions are shown in Figures S7-S12. The detachment forces are obtained from the force-extension profiles and are identically defined in both experiments and simulations (Figure S5, Figure S14).

The forced adsorption and desorption simulations were performed in the NAP_*z*_T ensemble. The temperature of the systems was fixed at 300 K using the velocity rescaling thermostat with stochastic term^57^ using a time constant of 1 ps. The pressure was kept at 1.0 bar using the Parinello-Rahman barostat^58^ with a time constant of 5 ps. Van-der-Waal interactions were cut-off by using a force-switch function between 1.0 to 1.2 nm. Coloumb interactions were calculated using the Particle Mesh Ewald (PME) method^59^ with a Fourier grid spacing of 0.12 nm. Bonds involving hydrogens were constrained using the LINCS algorithm^60^ with LINCS order 4. Periodic boundary conditions were applied in all directions and a time step of 2.0 fs was used. The simulations were performed with the Gromacs simulation package (versions 2018, 2021.5).^56^ Analysis was done using Gromacs and MDAnalysis python package^61^ and VMD^62^ was used for visualization.

#### Forces at different pulling rates

In dynamic pulling, the DNA was moved away from the mica surface with constant velocity. Eventually, the applied force led to the desorption of the DNA from the surface. The corresponding detachment force was obtained directly from experiments and simulations. By repeating such pulling simulations and experiments for a sufficient number of times one can obtain a distribution of the detachment forces including the mean detachment force and the most probable detachment force.

The dependence of the detachment force on the pulling velocity can be derived analytically assuming that the system diffuses on a free energy surface with a single sharp barrier and is pulled by a harmonic spring with constant velocity.^45,63^ For this current work, two regimes are of particular interest: For low pulling velocities *v* in the range of our experiments, the average detachment forces 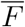 are expected to scale linearly with the logarithm of the force-loading rate ln(*k*_*s*_*v*).^64^ In this regime, the peaks in the force spectra correspond to the breaking of a single or a few molecular bonds and the detachment force is therefore largely independent of the length of the peptide chain. For very high pulling speeds in the range of our simulations, 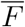 shows a different scaling behavior and is expected to scale with the square root of the loading rate 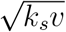.^65^

To test whether the predicted scaling behavior for simple model systems holds for the desorption of DNA at mica surfaces, we performed experiments and simulations at different pulling velocities. In addition to the simulations at *v* = 10, 1, 0.1 m/s, we performed two pulling simulations at 0.01 m/s and one at 0.005 m/s for systems with Na^+^. The loading rate was calculated from *k*_*s*_*v*, where *k*_*s*_ is the harmonic spring constant used in the simulations. Note that, for the simulation force-extension profiles, this loading rate is same as evaluated from the slope of force-time curve before detachment as done in experiments (Figure S14 B).

According experiments were performed at four different pulling velocities (0.01, 0.1, 1.0, 10.0 *µ*ms^*−*1^).

### Experiments

#### RCA Cleaning

The glassware and tweezers were immersed in a Radio Corporation of America (RCA) solution, 5 : 1 : 1, ultrapure H_2_O (18.2 MΩcm, Purelab Chorus 1, Elga LabH_2_O, Celle, Germany), NH_3_ (28.0-30.0%, Roth, Karlsruhe, Germany), H_2_O_2_ (*≥*30%, Sigma-Aldrich, St. Louis, MO, USA) at 70^*◦*^C for 40 minutes. Afterwards, they were rinsed three times with H_2_O, dried and stored at 120^*◦*^C.

#### Polymers

The DNA used for the AFM experiments was purchased as poly(T) single-stranded DNA (poly-T ssDNA, biomers, Ulm, Germany) with a thiol functionalized end group at 5^*′*^.

The DNA is constituted of 100 T nucleotides and thus, the contour length was calculated to be 67 nm based on the length of a base in ssDNA of 0.67 nm.^66^A triethoxysilane-polyethylene glycol-maleimide linker (silane-PEG-mal, Mw=5 kDa, NANOCS, Boston, MA, USA), with a contour length of approx. 41 nm was used to bind the DNA to the AFM cantilever tip.^67,68^

#### AFM Cantilever Tip Functionalization

Si_3_N_4_ AFM cantilevers, namely, MLCT-BIO-DC (force constants: 0.01 N/m −0.1 N/m, Bruker AFM probes, Camarillo, CA, USA) were used for all measurements. First, the cantilevers were activated with oxygen plasma (40%, 0.1 mbar, 2.0 min, Diener Electronics, Ebhausen, Germany) to gain hydroxyl groups on the surface of the cantilever tips. As a next step, a 5 kDa silane-PEG-mal linker was bound to the cantilever tip. The linker enabled us to couple a probe molecule to the cantilever tip via a covalent bond. Therefore, the cantilevers were incubated in a solution of silane-PEG-mal in toluene (99.99%, Fisher Chemicals, Hampton, NH, USA) at a concentration of 1.25 *mg/mL* for 2 h at approx. 22^*◦*^C. The cantilevers were rinsed in toluene, ethanol (*>*99.95%, VWR, France) and ultrapure H_2_O and finally incubated in a solution of poly-T ssDNA (100 nM) in 4-(2-hydroxyethyl)-1-piperazineethanesulfonic acid (HEPES) buffer (10 mM HEPES, NaCl 50 mM, pH 7, Sigma-Aldrich, St Louis, MO, USA) for 1 h at approx. 22 ^*◦*^C. Finally, the cantilevers were rinsed and stored in HEPES buffer at 4^*◦*^C.

Even though the cantilever tips are covered with maleimide groups, these undergo a hydrolysis (inactive PEGs) leaving just few binding sites for the single probe polymer to be attached. These inactive PEGs serve as a passivation layer to reduce undesirable interaction between the whole cantilever tip and an underlying mica surface as well as between a single PEG molecule and the cantilever tip. Furthermore, the choice of ssDNA length allows us to observe single ssDNA detachment events at a larger extension than the range of any remaining undesirable unspecific adhesion.

For every functionalization, control cantilevers were additionally prepared by the same procedure incubating the cantilevers in the pure solvent (HEPES buffer) instead of the poly-T ssDNA solution.

#### AFM Measurements

All measurements were performed on a MFP-3D-Bio and a Cypher ES (Asylum Research, an Oxford Instruments company, Santa Barbara, CA, USA) at approx. 298 K. A mica substrate (muscovite, diameter 10 mm to 12 mm, Plano, Wetzlar, Germany) was glued into a sample holder using a UV-curable adhesive (NOA 63, Norland Products, USA) and carefully cleaved with a clean scalpel until a flat surface was obtained. The measurement of poly-T ssDNA on mica took place in ultrapure H_2_O, LiCl, NaCl, KCl, CsCl, CaCl_2_ and MgCl_2_ (all 150 mM, Sigma-Aldrich, St. Louis, MO, USA). Before every measurement, the inverse optical lever sensitivity (InvOLS) was determined by fitting a linear function to the repulsive regime of a deflection-distance curve. In order to reduce errors, the determination of the InvOLS was performed by using an average of at least five individual InvOLS values. The spring constant of the cantilever was determined via the thermal noise method.^68,69^

The measurement parameters were defined as follows: force distance: 1 *µm*; pulling velocity for NaCl: 0.01 *µ*ms^*−*1^, 0.1 *µ*ms^*−*1^, 1 *µ*ms^*−*1^ and 10 *µ*ms^*−*1^, pulling velocity for all other experiments: 1.00 *µ*ms^*−*1^; contact force: 500 pN; sampling rate: 0.5-16.7 kHz; contact time towards the surface: 0-1 s. Force-extension curves were recorded in a grid-like manner with 10*·* 10 points covering 20*·* 20 *µm*^2^ (force maps) as well as in the continuous mode at different surface spots. At least two force maps were obtained per cantilever, pulling velocity and contact time towards the surface. First, control cantilevers carrying only a PEG functionalization were measured on mica in H_2_O (speific PEG-mica interaction observable, see Figure S13 A), then DNA functionalized cantilevers were measured on mica in H_2_O (strongly reduced DNA-mica interaction, see Figure S13 B), followed by measurements on mica in different ion chloride solutions, respectively.

For data evaluation, a self-programmed evaluation software based on Igor Pro (Wavemetrics, Portland, OR, USA) was used. The force-extension curves were corrected for drift^68^ and classified according to their shapes into plateau and stretching (including multiple stretches).^39^ The lower detection limit, due to the noise level of our instrument at the measurement conditions, allows us to detect any detachment events above 10 pN. In each force-extension curve the last detachment event, either a plateau or a peak at the end of a stretching event, was taken to determine the detachment force. The presented data comprise measurements with one to five different cantilever chips for each ion chloride solution.

For force loading rate dependent experiments with NaCl, stretching events were used. Using force-time plots, the loading rate was obtained for each single molecule detachment event. Due to the compliance of the PEG linker, the force loading rate could not be obtained from *k*_*s*_*v* as done for the simulations. Here, the slope of each force peak was determined by fitting a linear function in the force-time plot (Figure S14). The data points were divided into four different bins according to the different applied pulling velocities and the mean detachment force was evaluated for each bin. The total number of stretch curves vs all curves taken for the force loading rate dependent experiments with NaCl is: 78/1200 (7%, 0.01 *µ*ms^*−*1^), 280/600 (47%, 0.1 *µ*ms^*−*1^), 109/600 (18%, 1.0 *µ*ms^*−*1^), 122/1300 (9%, 10.0 *µ*ms^*−*1^).

## Results and Discussions

### Ion-specific interfacial structure

The negative surface charge of mica leads to the formation of a pronounced ionic double layer. In addition to the diffusive ions, which are loosely associated with the surface, the first layer of cations adsorbs specifically at the mica/water interface. Depending on the cation type, these ions occupy different binding sites: The ditrigonal cavities or the negatively charged oxygens bridging between the aluminum and silicon atoms (Figure 1 A).

The different binding modes are reflected in the position of the peaks in the ionic density profiles (Figure 1 C-H). The dominant peak in the Li^+^ density profile corresponds to the preferential binding at an oxygen triad above an Al atom. The two peaks of similar magnitude for Na^+^ reflect that Na^+^ binds to ditrigonal cavities and bridging oxygens. The single peak in the profiles for K^+^ and Cs^+^ reflects exclusive binding to the ditrigonal cavities. With the exception of Li^+^, the cation affinity on mica increases with decreasing ion size as reflected in the different number of cations needed to obtain a bulk salt concentration of 150 mM: Na^+^ ≤ Li^+^ < K^+^ < Cs^+^ (see methods). The affinity is also reflected in the resulting effective surface charge (Table 1).

For the divalent ions, Ca^2+^ binds to the oxygens bridging between aluminum and silicon. Mg^2+^ binds to the oxygens bridging between silicon atoms or between aluminum and silicon. Still, the affinity and the resulting effective surface charge are similar: Ca^2+^ ≤ Mg^2+^ (Table 1).

As a consequence of the different distribution and hydration of the cations, the density profile of the interfacial water shows pronounced differences (Figure 1 C-H). The strongly hydrated cations Li^+^, Mg^2+^, and Ca^2+^ cause a more structured and closer packed hydration layer compared to the other ions (Figure 1 C, G-H). Moreover, due to its strong hydration, Mg^2+^ binds in outer sphere configuration such that the interactions between Mg^2+^ and mica are always mediated by water.

The distribution of the cations at the mica/water interface and the structure of the interfacial water depends on the type of cation in the solution. Specific cation binding to the ditrigonal cavities and/or the bridging oxygen atoms compensates (Li^+^ and Na^+^) or even over-compensates (K^+^, Cs^+^, Mg^2+^, and Ca^2+^) the bare negative surface charge of mica (see Table 1). The resulting positive effective surface charge of mica leads to a long-ranged electro-static attraction of DNA or other negatively charged solutes. In addition, as evident from Figure 1B, the surfaced adsorbed cations form inner-sphere contacts with the phosphate oxygens of the DNA backbone. In summary, the subtle interplay of ion-specific adsorption, hydration effects, and the formation of direct cation-DNA contacts causes DNA to adsorb at negatively charged mica in the presence of mono- and divalent ions.

**Figure 1.**
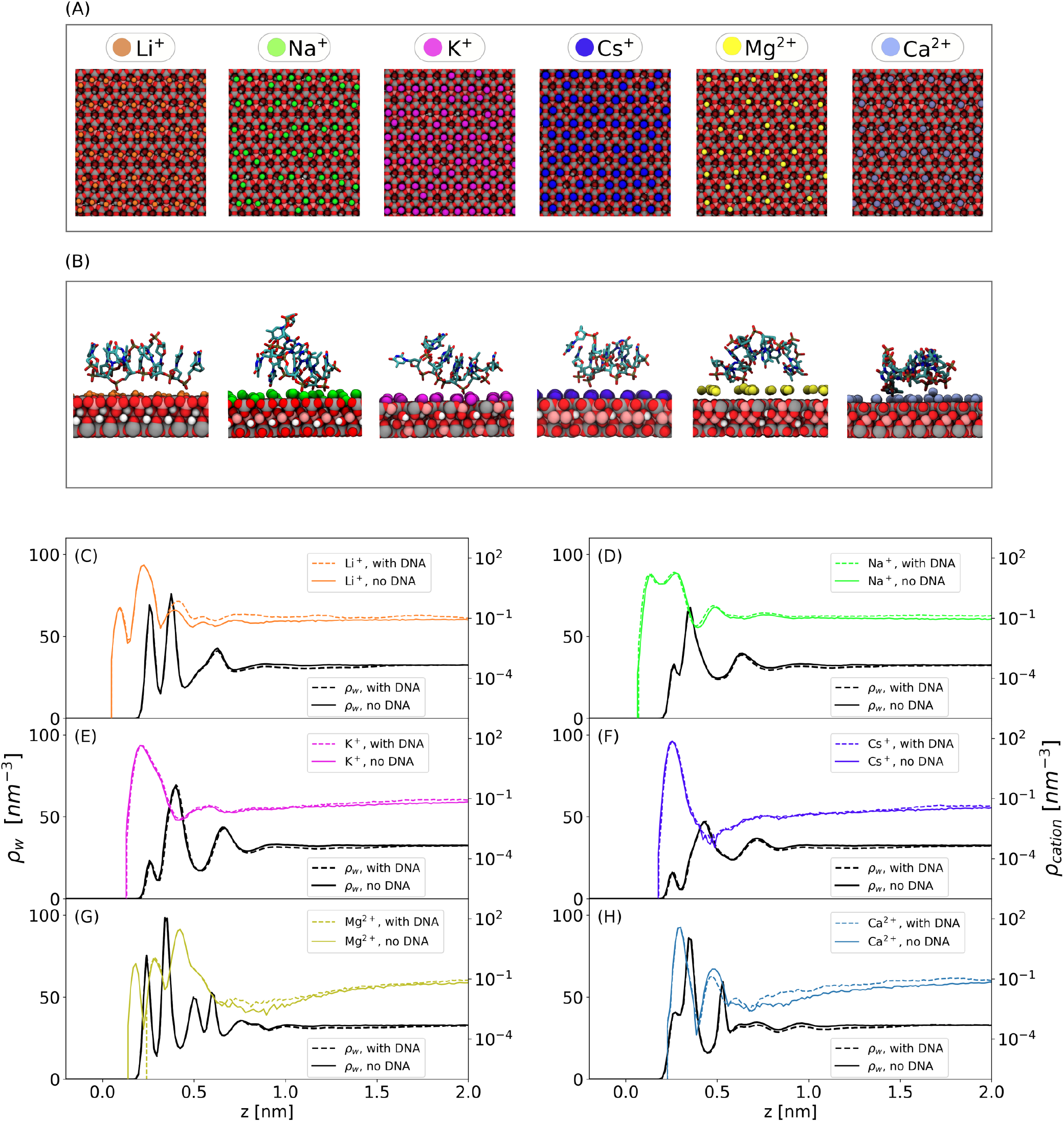
Distribution of cations, water, and DNA at the mica/water interface. (A) Simulation snapshots of the specifically adsorbed cations at the mica surface. The ion binding sites shift with increasing charge density from an oxygen triad to the center of a ditrigonal cavity. Silicon atoms are shown in gray, aluminum atoms in pink, and bridging oxygens in red. Water is not shown for clarity. (B) Simulation snapshots of the ssDNA at the mica/water interface for the different cations. (C-H) Ion distribution profiles and density profiles of the water oxygens. The profiles are shown in the absence (solid lines) and presence of the ssDNA (dashed lines). The bulk ion concentration in all cases is 150 mM. Anions are not shown. Similar locations of the density peaks perpendicular to the surface and the ion binding sites have also been noted previously^70,71^.

### Detachment forces in experiments and simulations

We now turn to the dynamics of DNA desorption from mica in NaCl solutions. In the dynamic pulling experiments and simulations, mechanical stress is built up via the external force. Eventually, the force leads to the desorption of the DNA from the surface. The corresponding force at detachment in dependence of the loading rate obtained from AFM experiments is shown in Figure 2A.

**Figure 2.**
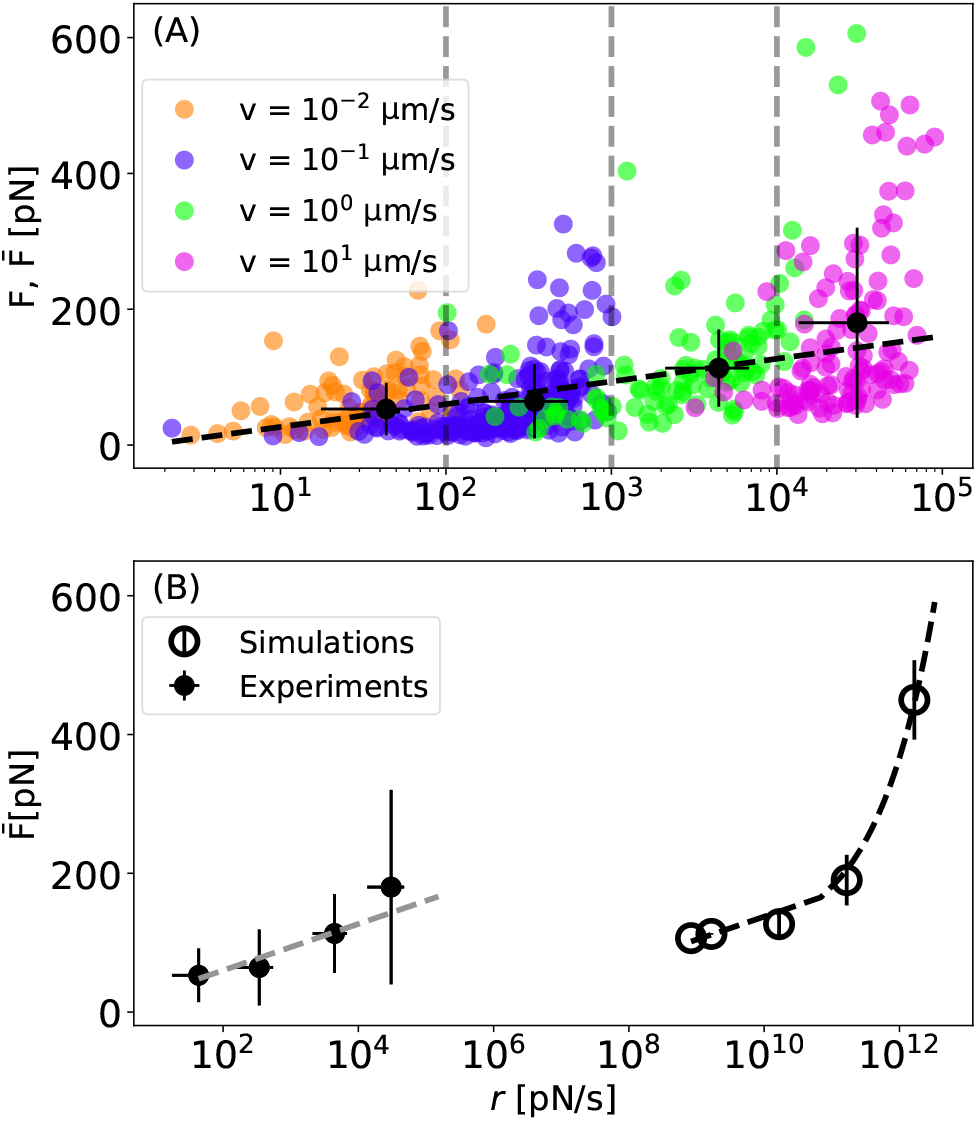
Dynamics of DNA desorption from mica in 150 mM NaCl. (A) Detachment force *F* and average detachment force 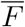 as a function of the loading rate for stretching events in 150 mM NaCl. Different colors indicate the four different experimentally applied pulling velocities. The average detachment force 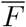 for blocking the loading rate into four bins is shown as black points. The error bars indicate the standard deviation. The black dashed line indicates a fit based on the Bell-Evan model.^64^ (B) Average force at detachment 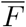 as a function of the loading rate *r* from experiments and simulations. The dashed lines correspond to fits according to the Bell-Evens model for experiments at low loading rates (gray) and the scaling with 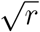 at high *r* observed in the simulations (black).

For the low pulling velocities, the average detachment force scales linearly with the logarithm of the loading rate. Here, we use the Bell-Evans model,^63,64^ wherein the bound state corresponds to the free energy minimum. The barrier separating the bound and the unbound state represents the transition state. The Bell-Evans model connects the average detachment forces to the loading rates with the intrinsic off rate *k*_0_ and the distance between bound and transition state *x*^*‡*^. We obtain *x*^*‡*^ = (0.28 *±*0.07) nm and *k*_0_ = (0.10 *±*0.12) s^*−*1^. These values indicate that the desorption process involves the breaking of strong local interactions instead of the gradual unbinding of the single-stranded DNA molecule as it is pulled away from the surface. For comparison, the measurement of polytryptophan and polytyrosin on a lipid bilayer showed a *x*^*‡*^ of 0.65 nm and 0.63 nm, respectively,^39^ unbinding of a FITC-E2 revealed a *x*^*‡*^ of about 0.4 nm^73^ and the rupture of a covalent siloxane bond revealed a *x*^*‡*^ of 0.021 nm.^74^ Note that the structure and charge of these systems, in particular for the lipid bilayer, differs significantly from the mica system investigated here.

A direct, quantitative comparison of the experiments and the simulations, as shown in Figure 2B, is challenging for two reasons: (i) The pulling velocities and loading rates in experiments and simulations differ by several orders of magnitude. With increasing velocities, the dissipative contributions due to friction increase and contribute significantly to the simulated detachment forces. (ii) The scaling deviates already at the low pulling velocities from ideal behavior (loading rate 4.8 *·*10^4^ pN/s in Figure 2 B). Deviations occur since the detachment force no longer corresponds to the breaking of a single contact between DNA and surface. The detachment forces then depend on the cantilever stifness and polymer length^75^ which differs by one order of magnitude in experiments and simulations. Hence, a simultaneous fit of experiments and simulations, established in uncharged systems,^39,44^ fails in these highly charged systems due to the strong interactions mediated by inner-sphere ion binding. Still, it is possible to compare the ion-specific contributions to the detachment forces from AFM experiments and simulations by using Na^+^ as a reference, as will be discussed further below.

### Comparison of the curve shapes

Comparison of the curve shapes and mobility of the DNA on the surface provides first insights into the desorption mechanism (Figure 3 and Table 1).

**Table 1:** Properties of the systems for different ion types. Effective surface charge density (*σ*_eff_) and diffusion coefficient *D* of DNA on the mica surface from simulations. Errors in *D* correspond to the standard deviation from block averaging. See supporting information section 1.2 and section 1.9 for details.

| Ion type | $\sigma_{\text{eff}}$ [ e/nm<sup>2</sup> ] | $D$ [ 10<sup>-7</sup> cm<sup>2</sup>s<sup>-1</sup> ] |
| --- | --- | --- |
| Li<sup>+</sup> | 0.13 | 0.65 ± 0.51 |
| Na<sup>+</sup> | 0.08 | 0.64 ± 0.20 |
| K<sup>+</sup> | 0.47 | 2.17 ± 0.61 |
| Cs<sup>+</sup> | 1.03 | 2.92 ± 1.15 |
| Mg<sup>2+</sup> | 0.51 | 5.32 ± 3.09 |
| Ca<sup>2+</sup> | 0.51 | 0.81 ± 0.16 |

**Figure 3.**
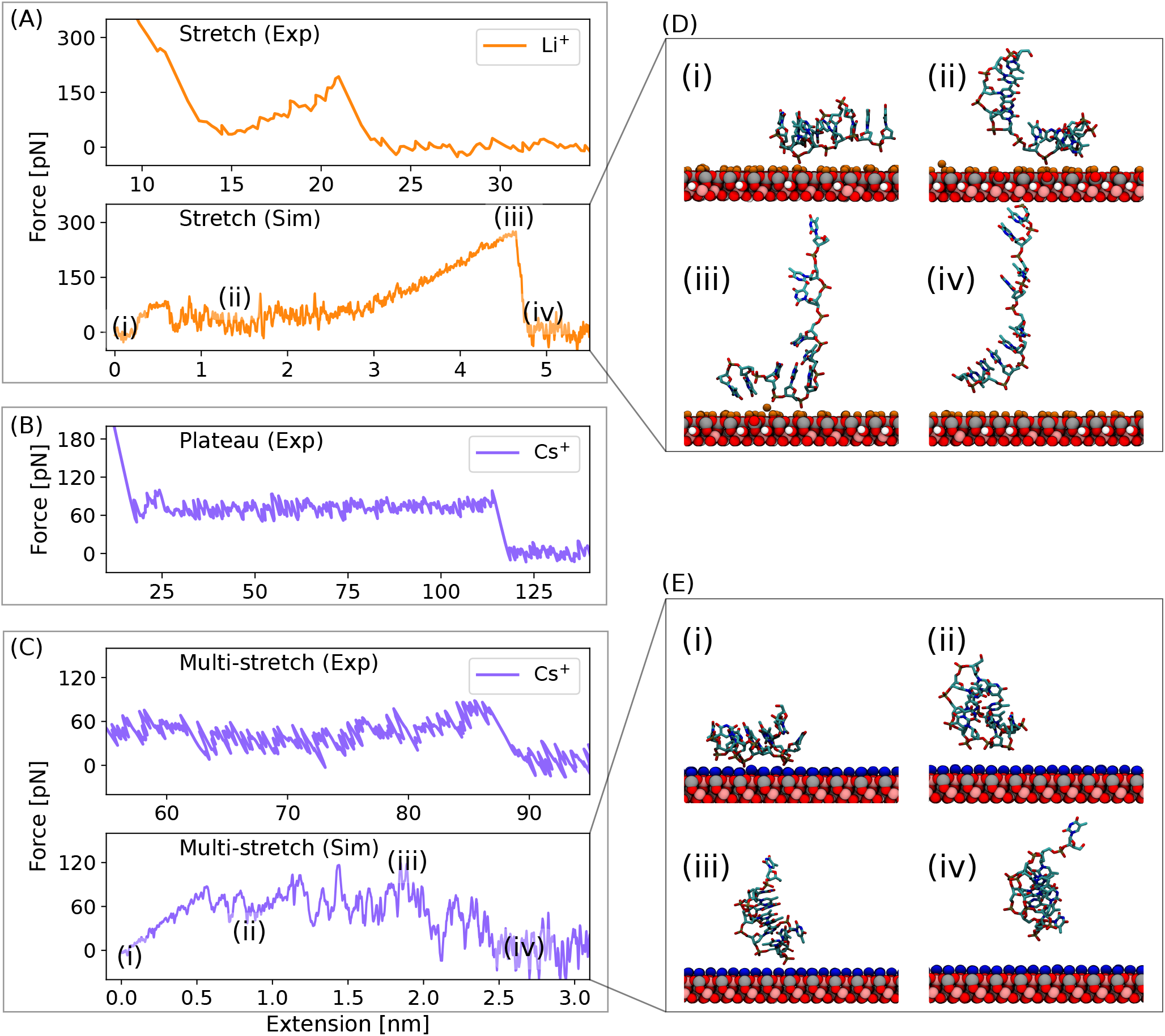
Comparison of the curve shapes. (A-C) Representative force-extension curves obtained from experiments and simulations for Li^+^ and Cs^+^. Note that plateau type curves are not observed in the simulations. High adhesion forces at low extension in the experimental curves result from unspecific adhesion peaks at small extensions due to an interaction of the whole AFM cantilever tip and the underlying mica surface. (D-E) Snapshots of the DNA at different points during the desorption process indicated in the corresponding simulated force-extension curves. Here, the force-extension curves correspond to pulling velocities of 0.1 m/s for simulations and 1.0 *µ*m/s experiments. The slope of the final drop of the force is related to the respective force constant of the cantilever.^72^

The different curve shapes indicate different desorption mechanisms and can be classified into different types (Table S1 and Figures S15 and S16): plateaus, stretching, and multiple stretches. Stretches are observed most frequently (Figure 3A,D and Table S1). Here, individual contacts are very stable and relax slowly. The DNA is stuck on the surface with low mobility and application of force leads to stretching of the DNA molecule. Flat force plateaus occur in the experiments (Figures 3B, S15 and S16) but are never observed in the simulations in agreement with previous results.^38,39^ Here, the DNA-surface interactions relax faster than the AFM cantilever tip is moved away from the surface. Due to the high pulling velocities, such a stationary non-equilibrium is not reached in the simulations. Finally, an intermediate situation corresponds to a stick-slip motion (Figure 3C). Here the DNA switches between surface adsorbed and desorbed conformations (Figure 3E). Furthermore, in the experiment, the already desorbed ssDNA strand could act as a flexible tether. The latter averages out the force acting on the AFM cantilever, possibly leading to flat force plateaus. In the simulations, the tether is much shorter leading rather to peaks in the force-extension response.^38,39,47,76^

The simulations provide further insights into the desorption mechanism: Convex stretching is predominately observed for cations with high charge density as in the example for Li^+^ (Figure 3A, D). The lifetime of a contact formed between the adsorbed cation and the DNA is on the order of 2 ns^34^ and remains stable while the DNA is stretched. At the same time, the strongly bound hydration layer prevents rebinding of the DNA and the force-extension curves show a single, well-pronounced detachment event. Ions with low charge density, such as K^+^ and Cs^+^, often show multiple stretches (Figure 3C, E) due to the shorter lifetime of contact pairs (about 0.1 ns) and the lower binding affinity.^34^ Here, the contacts often break before the DNA is stretched. Due to the weaker hydration layer, rebinding is frequently observed leading to the characteristic stick-slip motion. Note that in addition to the ion-mediated DNA-surface interactions, the conformational changes of the DNA (Figure 3D, E) influence the exact shape of the force-extension curves.

### Origin of ion-specific detachment forces

Figure 4 shows the detachment forces for all ions from simulations and experiments relative to the average force in NaCl. Overall, the distributions are broad indicating that different desorption mechanisms contribute to the average force. The variety of contributions is larger than the difference between the individual ions obscuring ion-specific effects. The overall trend in simulations and experiments is similar: Forces for Cs^+^ are typically lower compared to Na^+^, while K^+^ and Na^+^ are similar. For Li^+^, Ca^2+^, and Mg^2+^ the forces of individual events can be higher or lower compared to Na^+^.

To provide further insights into what contributes to high and low desorption forces, we classify the type of contact that breaks during detachment into four categories: (i) Water-mediated contacts: The water-mediated contact between DNA and the cation breaks (Figure 5A). (ii) Ion-phosphate oxygen contact: The direct contact between phosphate oxygen and ion breaks; the ion remains on the surface (Figure 5B). (iii) Ion-surface contacts: The direct contact between ion and surface breaks; the ion leaves the surface with the DNA (Figure 5C). This class comprises the special case in which the contact between hydrated Mg^2+^ and the surface breaks (Figure 5C). (v) Ion-phosphate and ion-surface contacts: One ion-surface and one ion-phosphate oxygen contact break simultaneously. This occurs in a few cases when the non-bridging phosphate oxygen is coordinated by two surface-bound cations (Figure 5D). For each category, the force necessary to break the contact depends on the ion type. For water-mediated contacts and for ion-phosphate interactions, the forces increase with increasing charge density according to Cs^+^ *<* K^+^ *<* Na^+^ *<* Li^+^ *<* Ca^2+^ *<* Mg^2+^.^34,77^

The ion-surface forces reflect the binding affinity to mica with ordering Mg^2+^ *L*Ca^2+^ and Na^+^ *L*Li^+^ *<* K^+^ *<* Cs^+^. Due to the contributions of all contact types to the resulting detachment forces (Figure 5E-J), it is not surprising that the average desorption forces do not show a clear ion specific trend (Figure 4, Table S2). Rather, the detachment forces originate from multiple pathways due to the complex interplay of ion-DNA, ion-surface, and hydration interactions.

**Figure 4.**
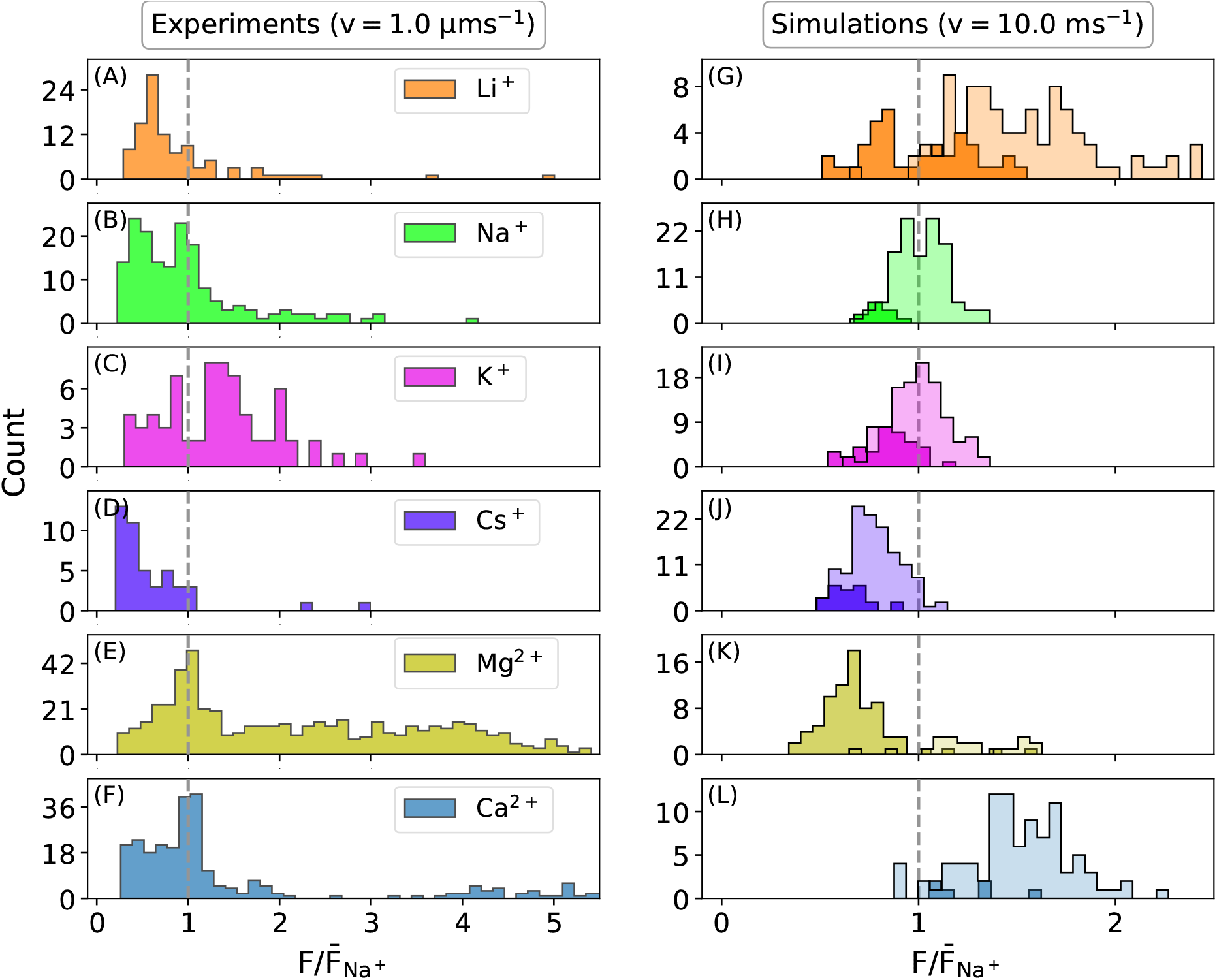
Distribution of detachment forces *F* from simulations and experiments with the average detachment force 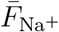 in NaCl as reference (indicated by vertical dashed lines). In the simulation data, the forces are further divided into two classes according to the desorption mechanism: Water-mediated contacts (dark shades) and direct ion contacts (light shades).

**Figure 5.**
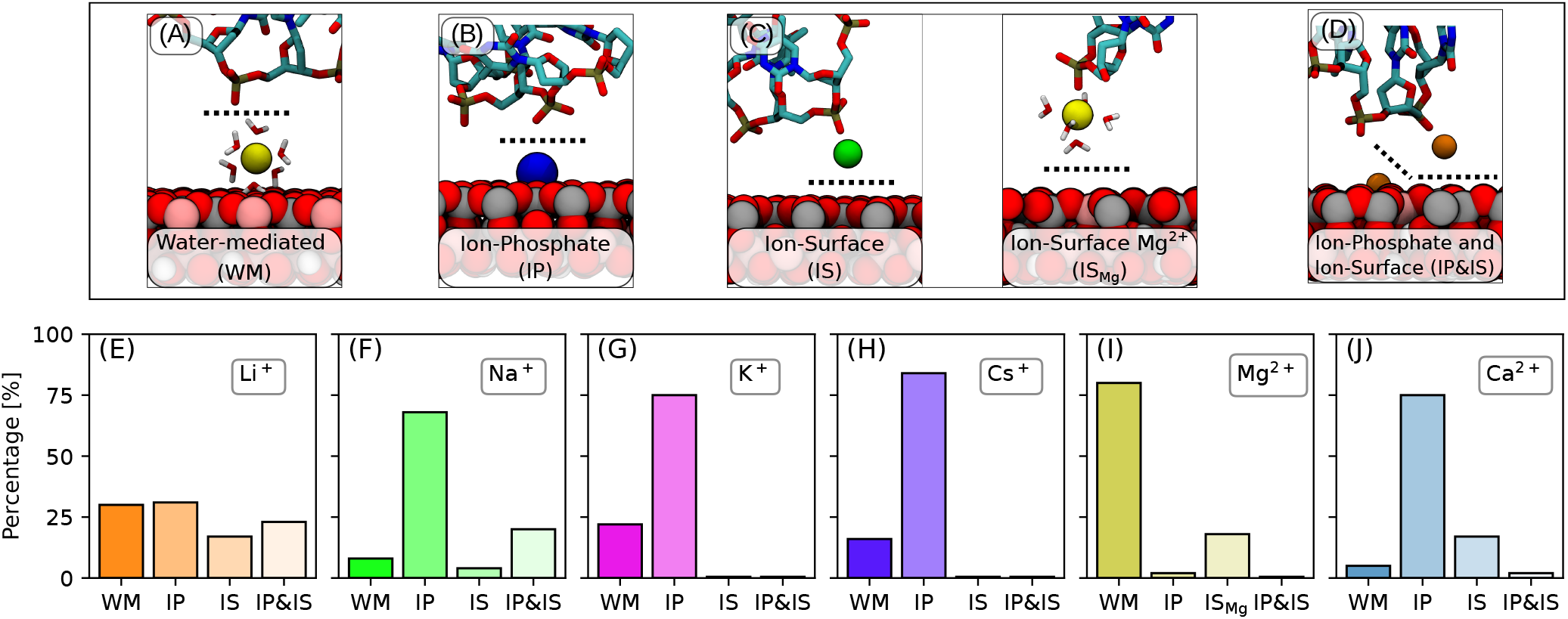
Classification of the type of contact breaking during detachment. Illustration of the four categories by simulation snapshots (A-D). The dashed lines show the point where contacts break. (E-J) Percentage of different contact types observed in the detachment forces for different ions.

Overall, the detachment forces induced by water-mediated contacts are lower compared to forces induced by breaking of direct ion-surface or ion-DNA contacts (Figure 4). The splitting into water-mediated and direct forces further elucidates the broad and sometimes bimodal distribution observed in the experiments. For example, Mg^2+^ has a pronounced peak in the distribution at detachment forces that are lower compared to the Na^+^ reference (Figure 4K). These result from water-mediated contacts which are predominant for Mg^2+^, but much less frequent for Na^+^ (Figure 5F, I). In addition to these low detachment forces, ion-surface mediated interactions contribute to a second mode of forces that are larger than the Na^+^ reference. However, due to the strong hydration of Mg^2+^ the outer-to-inner sphere transition is slow. ^53,78^ Using a high temperature pre-equilibration (see methods) allowed us to generate inner-sphere contacts which result in higher desorption forces and closer agreement with the experimentally observed high detachment forces (Figure S17).

The direct comparison of force distributions from experiments and simulation highlights the importance of accurate and balanced force fields but also reveals shortcomings of the current parameters. For instance, for Ca^2+^ the low force mode is absent. Clearly, the water-mediated pathways are underrepresented. This is likely caused by a too high surface affinity since the mica-Ca^2+^ interactions have not been optimized.^52^ By contrast, for Li^+^ the high detachment force is over-represented. This is likely caused by a too high binding affinity toward mica and/or the phosphate oxygen. To provide an improvement of the force field parameters, accurate experimental data for the binding affinities to mica is required to correctly balance the interactions with mica, DNA and water.^52,53^

## Conclusions

In summary, we characterized the detachment forces of single-stranded DNA at mica surfaces in the presence of different mono- and divalent cations from combined all-atom MD simulations and single-molecule AFM experiments. At the negatively charged mica surface a pronounced ionic double layer is formed. Different ion types occupy different binding sites and their binding affinity increases in the order Mg^2+^ *L*Ca^2+^ and Na^+^*L* Li^+^ *<* K^+^ *<* Cs^+^. Ion specific binding overcompensates the bare negative surface charge of mica. The effective surface charge is lowest for Na^+^ and largest for Cs^+^ and gives rise to long-ranged attraction. In addition, direct and water-mediated contacts are formed between the surface-adsorbed cations and the DNA. Formation of inner-sphere contacts requires partial ion dehydration and is hence most difficult for strongly hydrated ions. Once formed, the strength of such inner-sphere contacts increases with increasing charge density according to Cs^+^ *<* K^+^ *<* Na^+^ *<* Li^+^ *<* Ca^2+^ *<* Mg^2+^.^52^ As a consequence, DNA adsorption at mica results from a complex interplay of ion-surface, ion-DNA interactions as well as hydration effects. It is therefore not surprising that the detachment forces do not show a unique ion specific trend. Rather, the detachments forces for each ion type originate from multiple pathways: Detachment forces induced by water-mediated contacts are low while the breaking of direct ion-surface or ion-DNA contacts results in higher forces. The splitting into water-mediated and direct forces further elucidates the broad and sometimes bimodal distribution observed in the experiments.

We conclude that Cs^+^ results in the lowest detachment forces while the other monovalent ions yield similar results. For monovalent ions, K^+^ is the best choice due to the largest surface coverage. To immobilize DNA further, the physiologically relevant ions Ca^2+^ and Mg^2+^ are ideal to form a few specific and long-lived contacts between DNA and the mica surface in agreement with recent results.^5^

## Acknowledgements

This research was supported by the Bundesministerium für Bildung und Forschung (BMBF) within the Röntgen-Angström cluster project Medisoft, number 05K18EZA. The work was also supported by the Emmy Noether program of the Deutsche Forschungsgemeinschaft (DFG, German Research Foundation), number 315221747. We acknowledge the GOETHE HLR for super computing access. The authors gratefully acknowledge the scientific support and HPC resources provided by the Erlangen National High Performance Computing Center (NHR@FAU) of the Friedrich-Alexander-Universität Erlangen-Nürnberg (FAU) under the NHR project b119ee. We thank the Deutsche Forschungsgemeinschaft (DFG, German Research Foundation) under Germany’s Excellence Strategy -EXC-2193/1 390951807 (CW, ML and BNB).

## Supporting Information Available

Further force-extension curves from simulations and experiments, simulations and experimental protocols and analysis methods.

