## Supplementary Information for "Adsorbing DNA to mica by cations: Influence of the valency and ion type"

### 1 Simulation Details

#### 1.1 Obtaining the correct bulk salt concentration in simulations

To obtain a bulk salt concentrations $c$ in simulations box containing $N_w$ water molecules, the number of cations or anions, $N_{ion}$, required is given by the following simple relation,

$$N_{ion} = \frac{N_w c}{55.55} \tag{1}$$

where c is the molar concentration. If there is an additional charged component in the system like a DNA molecule, extra ions are added to neutralize the whole system. During the course of simulations, the oppositely charged ions condense around the DNA to compensate for the negative charge. However, in presence of highly charged surfaces like mica in our case, oppositely charged ions not only compensate for the surface negative charge, but overcompensate it. As there are a finite number of ions in our simulation system, this leads to a decrease in the bulk ion concentration. This decrement in bulk concentration depends on the affinity of a given ion toward the mica surface. Therefore, the amount of ions required to obtain a given bulk salt concentration depends on the type of ion.

Figure S1 A shows the cation concentration profile for $K^+$ ion with different added concentrations according to Eqn. 1. The obtained bulk concentration is smaller than the added concentration as shown in Figure S1 B. There is a linear relationship between added and obtained bulk concentration. The slope of this relationship varies among different ions based on their affinity towards the mica surface. In order to attain a bulk concentration of 150 mM for a specific ion, two simulations was performed at a higher and lower concentrations. By assuming a linear relation between the added and obtained concentrations, the required concentration to achieve 150 mM was interpolated.

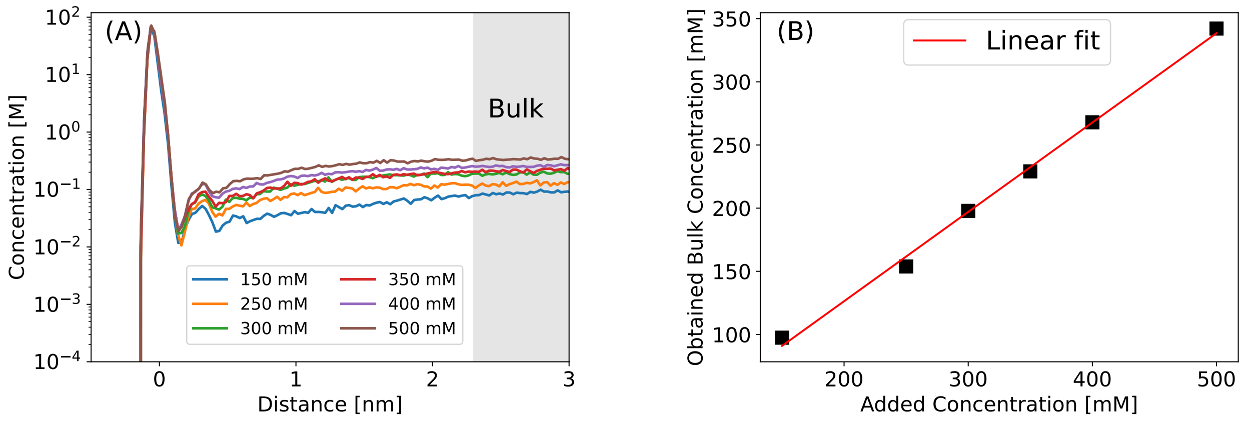


Figure S1: (A) Concentration profile for $K^+$ ion for different added concentrations according to eqn. 1. The added concentrations are shown in the legend. These profiles are obtained by considering the last 50 ns of a 100 ns equilibration run. (B) There is a linear relationship between the added concentration and the obtained bulk salt concentration.

#### 1.2 Effective surface charge density

The number of ions bound to the surface is obtained from the peak of the cation number density $\rho_n(z)$ according to:

$$N_{surf} = \int_0^z \rho_n(z') l_x l_y dz', \tag{2}$$

where, $l_x, l_y$ are the $x$ and $y$ dimensions of the mica surface. The upper limit $z$ is defined by the point where $N_{surf}$ almost saturates, because the ion density in the bulk is orders of magnitude less than in the adsorbed double-layer region (Figure S2 B). We obtain 136 $Li^+$, 133 $Na^+$, 156 $K^+$, 189 $Cs^+$, 79 $Mg^{2+}$ and 79 $Ca^{2+}$ adsorbed cations. Therefore, the adsorbed charge is given by $q_{adsorbed} = vN_{surf}e$, where $v$ is cation valency. The charge on the bare mica is $q_{bare} = -128e$. The effective surface charge density of mica is given by:

$$\sigma_{eff} = \frac{q_{adsorbed} + q_{bare}}{l_x l_y}. \tag{3}$$

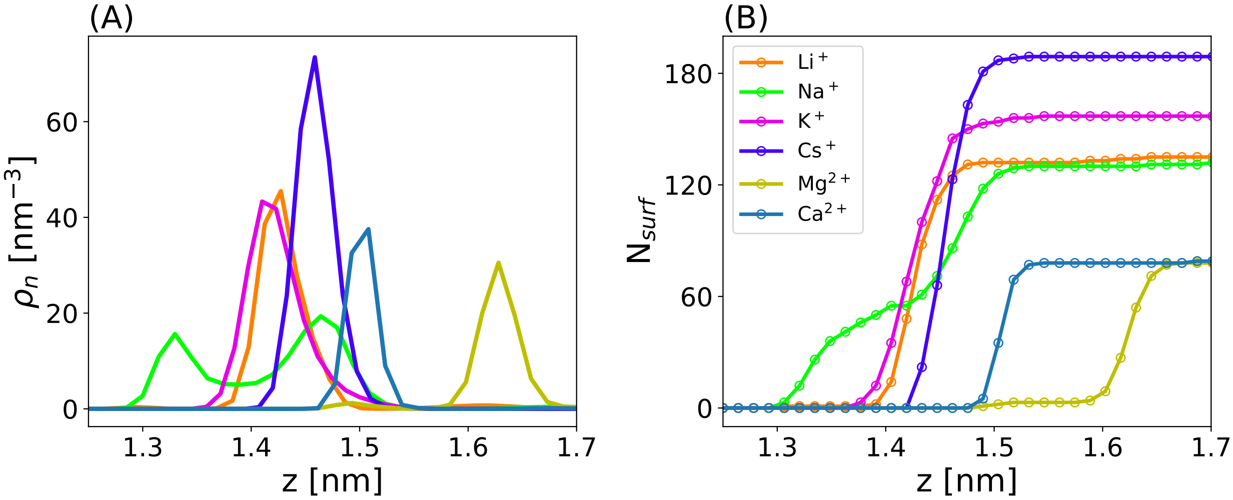


Figure S2: (A) Zoomed view of cation number density $\rho_n$ representing the adsorbed double-layer of ions. (B) The number of adsorbed ions obtained from (A) using eqn. 2.

#### 1.3 Constant force simulations

The DNA molecule, initially at 2 nm distance from the center of mass of mica surface atoms is moved towards the mica surface by applying an external force $\mathrm{F_{ext}} = \Sigma_\mathrm{i} \mathrm{m_i a}$, where $m_i$ is the mass of a heavy atom and $a$ is the acceleration. The acceleration is varied from 0 to 0.1 nm/ps$^2$ which corresponds to varying $\mathrm{F}_{ext}$ from 0 to 470 pN. For a given value of $\mathrm{F}_{ext}$, the DNA center of mass attains an equilibrium distance $\mathrm{z}_{eq}$. The amount of work done for bringing DNA from bulk to a distance of $\mathrm{z}_{eq}$ is given by:

$$W(z) = -\int_2^z F_{ext} dz'. \tag{4}$$

The total work of adhesion $A$ is given by $A = W(z_{eq})$. The evolution of $z$ and $W$ for $\mathrm{F}_{ext} =$ 470 pN is shown in Figure S3 (A,B).

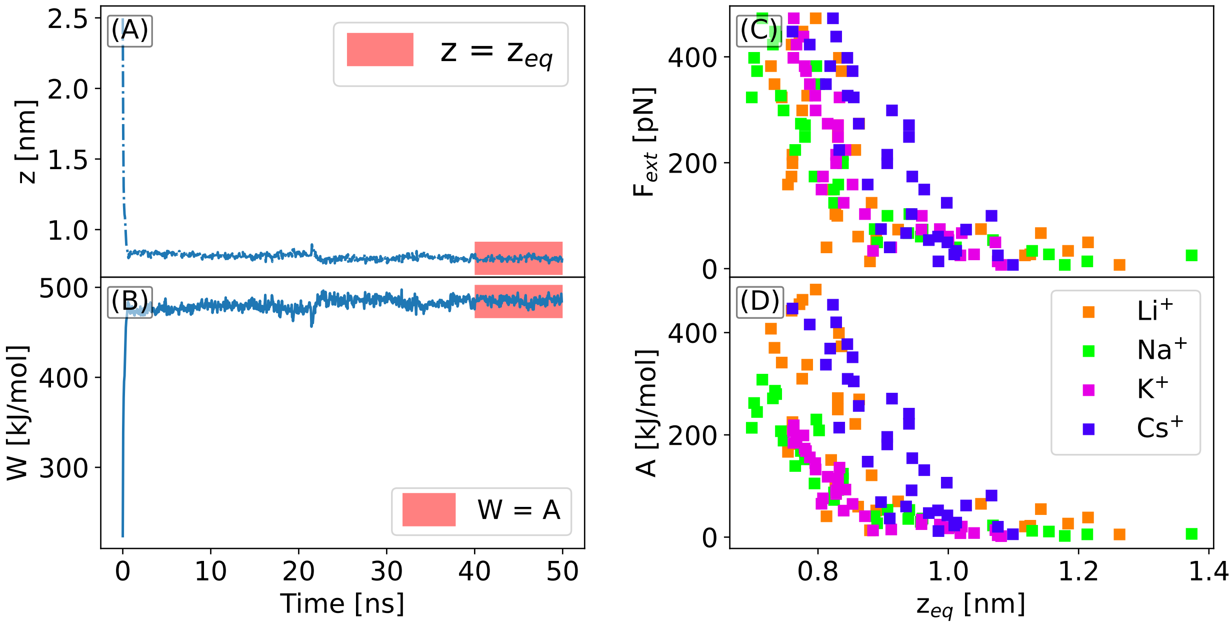


Figure S3: (A) DNA center of mass $z-$distance from a mica surface for a constant external force $F_{ext} = 470$ pN. (B) Work done evaluated using eqn. 4. The average of z and W in the red-shaded region is assumed to be $z_{eq}$ and work of adhesion $A$. (C) $F_{ext}$ vs $z_{eq}$. (D) A vs $z_{eq}$.

#### 1.4 Distance of each DNA residue from the mica surface during equilibration

After the forced adsorption, before pulling, we allow the DNA to relax and obtain an equilibrium conformation on the mica surface. The conformation is monitored by looking at the minimum $z$ distance of the phosphate group of each residue from the mica surface (Figure S4)

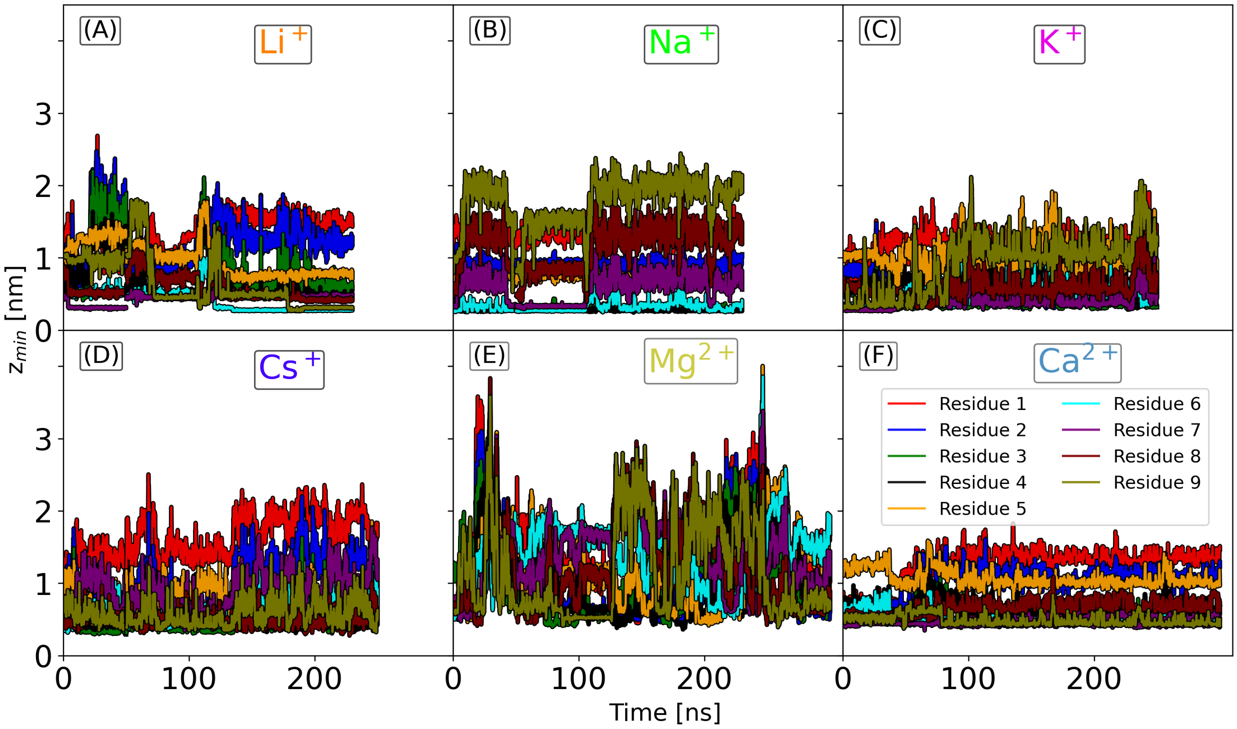


Figure S4: Equilibration after forced adsorption with $F_{ext}$ = 470 pN. The plots show the evolution of the minimum distance of the DNA phosphate group of each residue from the mica surface. Configurations equally spaced in time after 100 ns or 150 ns are selected for pulling.

#### 1.5 Definition of detachment force

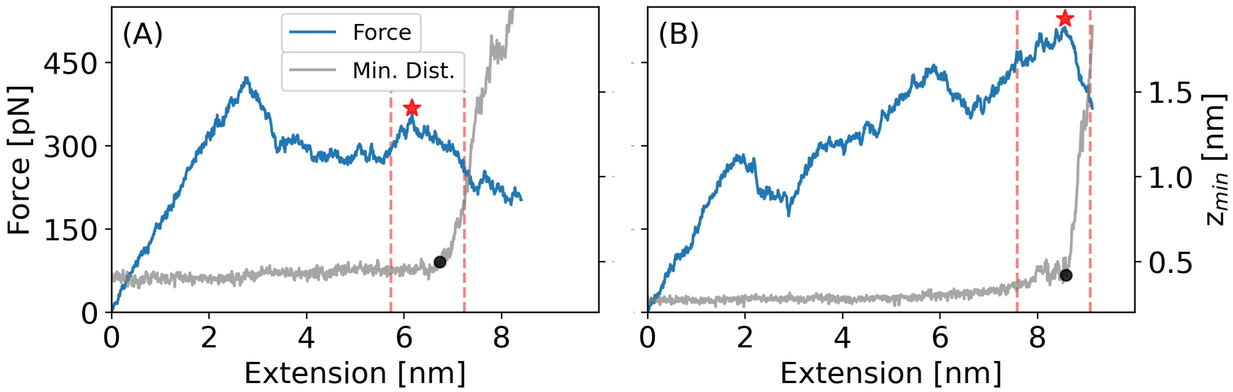


Figure S5: To be consistent we use the same definition of detachment force as in the experiments (Figure S14): The minimum distance (Min. Dist.) of DNA from the mica surface is monitored during the pulling of DNA. Min. Dist increases sharply at the desorption point indicated by the black circular marker. The maximum force within 1.5 nm of the desorption point is taken as the force at the detachment indicated by the red star marker. (A) In this case, the force at detachment is smaller than the maximum force over the course of desorption process (over the whole force-extension curve). (B) Here, the detachment force is the same as the maximum force over the course of desorption process. These plots corresponds to a pulling velocity of 10 m/s.

#### 1.6 Analysis of type of contacts that break at rupture

The analysis of interactions that break at rupture (main text Figure 5 (E-J)) is done using a custom python script based on MDAnalysis.[1] The script does the following:

1. Identify the detachment point in the force-extension profile. The detachment point is detected from the minimum distance of DNA from mica shown by the gray curve in Figure S5. At detachment the minimum distance increases sharply as shown by the black circular marker in Figure S5 A, B.

2. Obtain the frames of interest (FOI) in the simulation trajectory that are within ∼1 nm on either side of the detachment point in the force-extension curve.

3. For the FOI, three cation lists are created: (i) L1: those within a cut-off distance ($r_1$) of the non-bridging oxygen of the phosphate backbone, i.e., O1P and O2P, (ii)

L2: Those within a cut-off distance ($r_2$) of the DNA nucleobase oxygens, i.e., O2 and O4, (iii) L3: Those within a cut-off distance ($r_3$) of mica surface oxygen atoms. The pair lists are created using the MDAnalysis `capped_distance` function. The cut-off distances ($r_1, r_2, r_3$) are determined from the radial distribution function (RDF) of the cation and the water oxygen and are shown as vertical dashed lines in Figure S6. For all ions except $Mg^{2+}$ $r_1 = r_2 = r_3$. For $Mg^{2+}$, since all the surface-bound $Mg^{2+}$ is in hydrated state, $r_3$ is set by the second RDF maximum (Figure S6 E), while $r_1 = r_2$ is the same as for other ions.

4. The number of shared cation between the phosphate oxygens and the mica surface ($N_p$) is given by the intersection of L1 and L3, while the number of shared cation between the DNA nucleobase oxygen and the mica surface ($N_n$) is given by the intersection of L2 and L3.

5. If $N_p = N_n = 0$: There are no shared cations at any point during the detachment process. Therefore, the rupture force is assumed to arise from indirect water-mediated contacts.

6. If $N_p \neq 0$ or $N_n \neq 0$: We look at the last FOI, where the DNA is fully detached from the mica surface. We count the number of initially shared cations that are still bound to the mica surface. For $n_p$ out of $N_p$ and $n_n$ out of $N_n$ cations bound to the surface, it follows:

   (1) If $n_p$=$N_p$: All shared cations are bound to the mica surface and the rupture force is attributed to the breakage of the ion-phosphate (IP) type of contact.

   (2) If $n_p = 0$: All shared cations are desorbed from the mica surface. Therefore, the rupture force is attributed to breakage of the ion-surface (IS) type of contact.

   (3) If $n_p \neq 0$ and $n_p < N_p$: Some shared cations are bound to the mica surface and some are bound to the DNA phosphate. Therefore, the rupture force arises from simultaneous breakage of both ion-phosphate (IP) and ion-surface (IS) type of contacts.

In almost all cases for all types of ions, we find that $N_n$ is close to zero, i.e., the nucleobase is not involved in the interaction. Therefore, the classification for ion-nucleobase is omitted.

The visual inspection (in VMD) of random trajectories ($> 10$) accurately matches the results from the script given above.

#### 1.7 First hydration shell of mono- and divalent cations

The cation-water radial distribution functions provide insights into the stability of the first hydration shell. The height of the first peak reflects the ion-water interaction strength and follows the order $Cs^+ < K^+ < Na^+ < Li^+ < Ca^{2+} < Mg^{2+}$ (Figure S6) in agreement with the experimentally measured hydration free energies.[2] The position of the first peak corresponds to the radius of the hydration shell. Moreover, since the radial distribution function is connected to the free energy via Boltzmann inversion, the value -$k_B T \ln g(r_{\min})$ at the minimum corresponds to the free energy barrier of water exchange. Smaller barriers lead to faster exchange and a less robust first hydration shell. For the ions investigated here, the exchange is fastest for $Cs^+$ and slowest for $Mg^{2+}$ and otherwise follows the order: $Cs^+ > K^+ > Na^+ > Li^+ > Ca^{2+} > Mg^{2+}$ (Figure S6) in agreement with experimental results.[3]

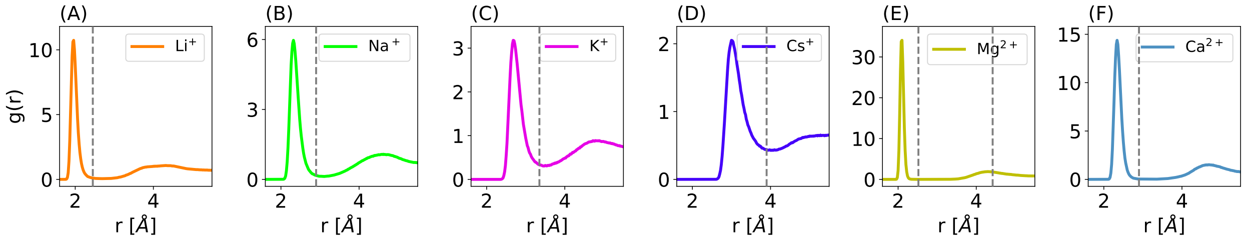


Figure S6: Radial distribution function, $g(r)$, between water oxygens and the cations. The vertical dashed lines represent the cut-off position used to define direct and indirect contacts while assigning the type of contacts that break at rupture.

#### 1.8 Force-extension profiles from simulations at low pulling speed (0.1 m/s)

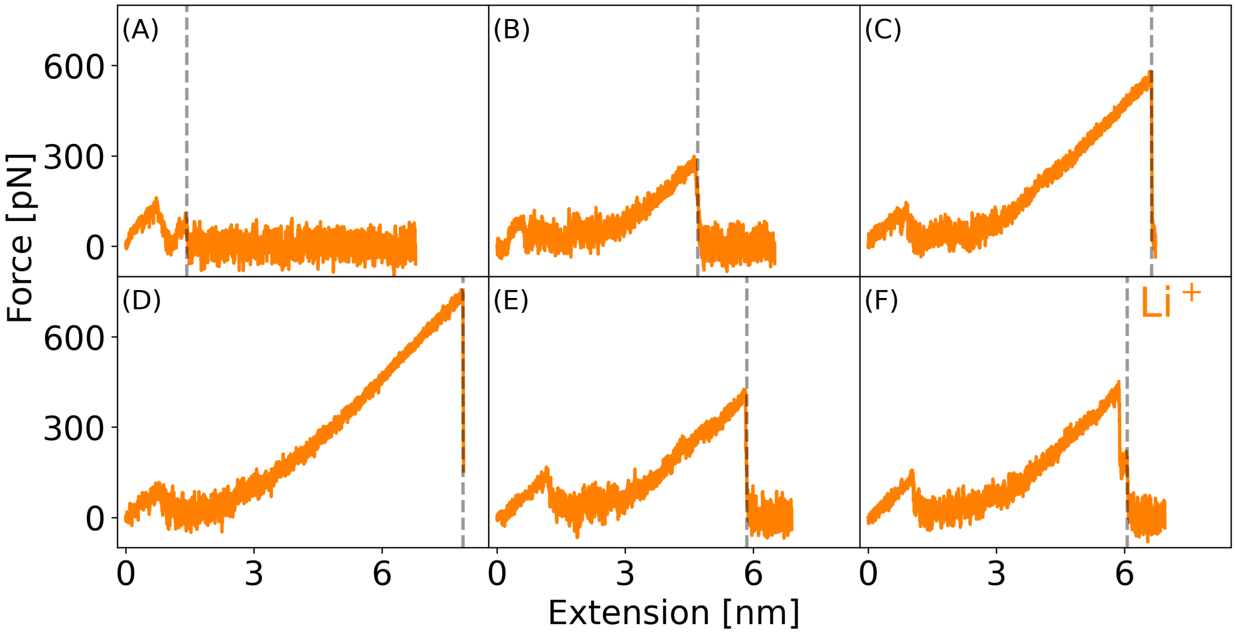


Figure S7: Force-extension profiles from simulations for $Li^+$ at a low simulation pulling speed of 0.1 m/s. The dashed vertical lines show the point where the DNA fully detaches from the mica surface.

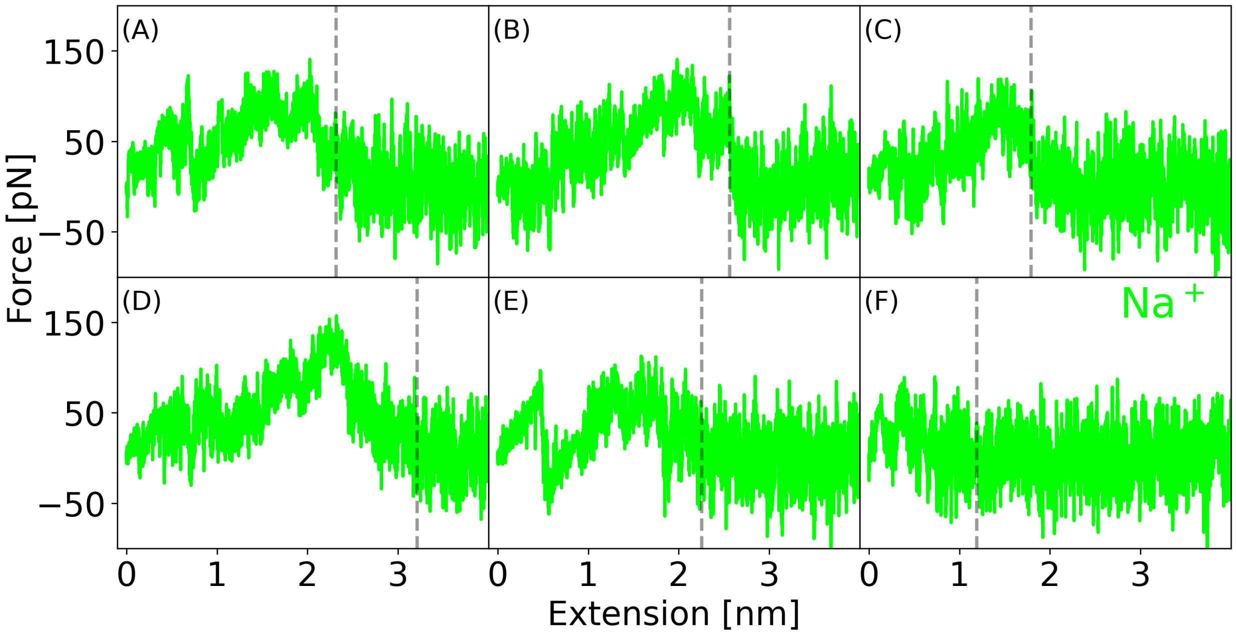


Figure S8: Force-extension profiles from simulations for $Na^+$ at a low simulation pulling speed of 0.1 m/s. The dashed vertical lines show the point where the DNA fully detaches from the mica surface.

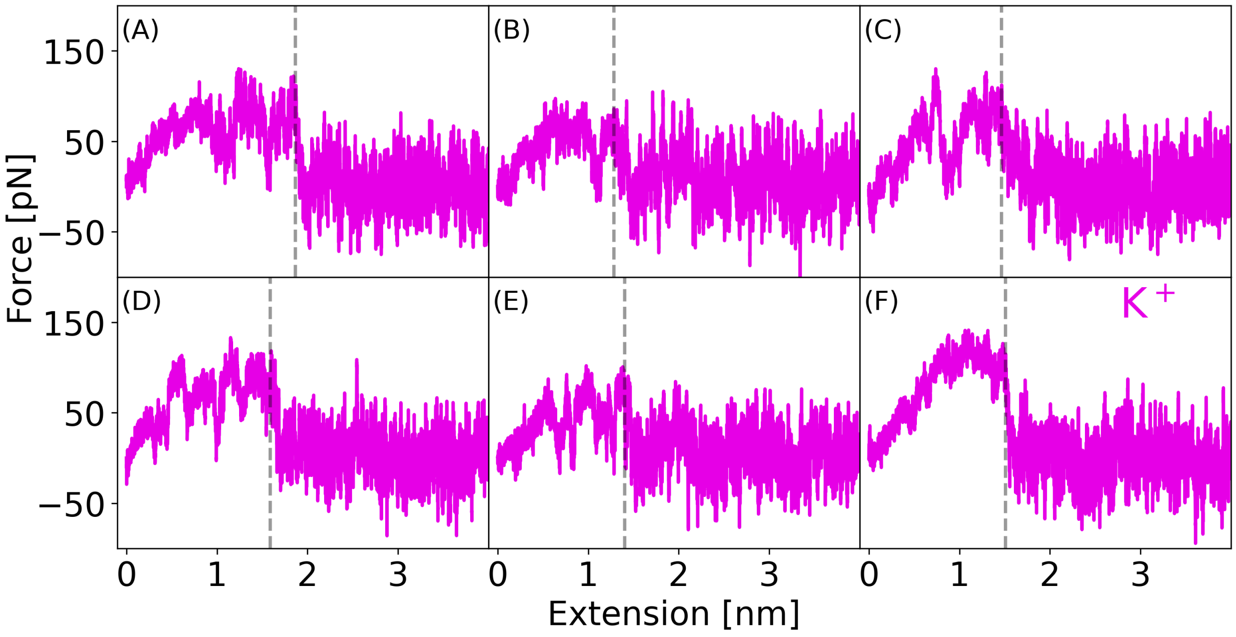


Figure S9: Force-extension profiles from simulations for $K^+$ at a low simulation pulling speed of 0.1 m/s. The dashed vertical lines show the point where the DNA fully detaches from the mica surface.

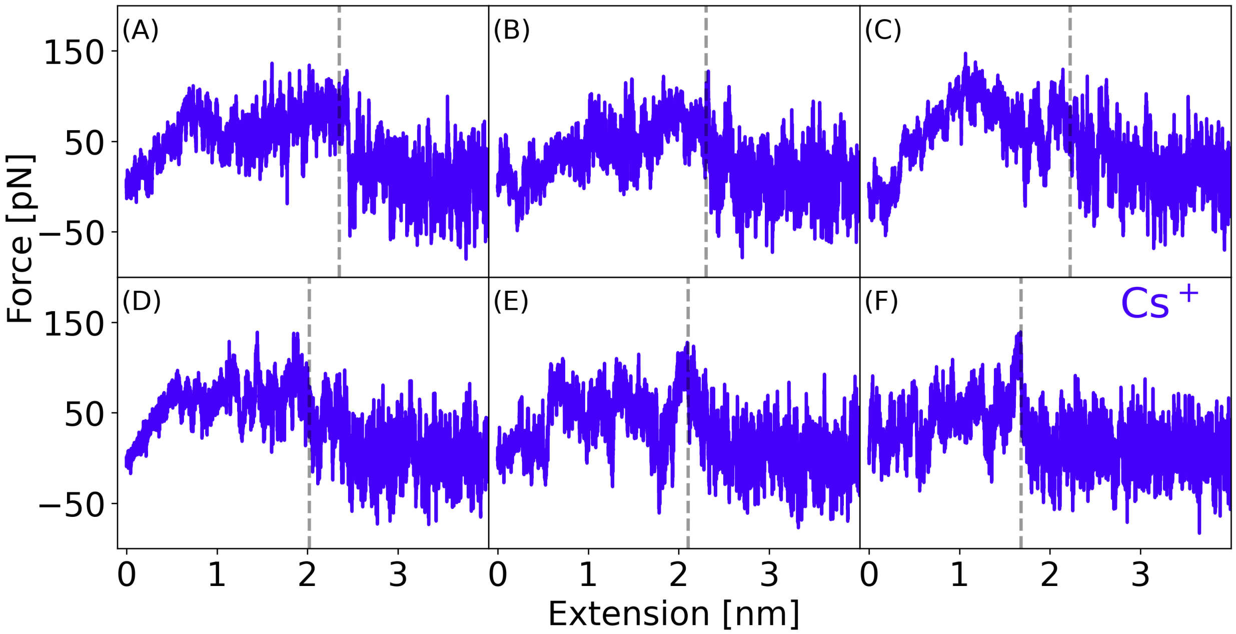


Figure S10: Force-extension profiles from simulations for $Cs^+$ at a low simulation pulling speed of 0.1 m/s. The dashed vertical lines show the point where the DNA fully detaches from the mica surface.

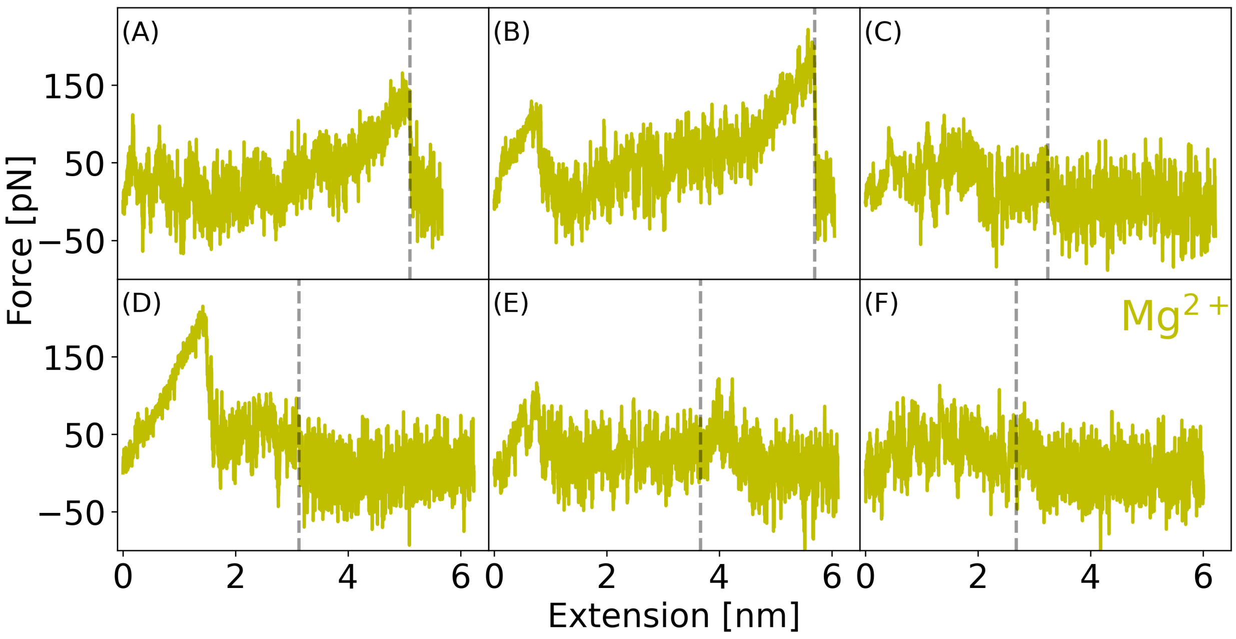


Figure S11: Force-extension profiles from simulations for $Mg^{2+}$ at a low simulation pulling speed of 0.1 m/s. The dashed vertical lines show the point where the DNA fully detaches from the mica surface.

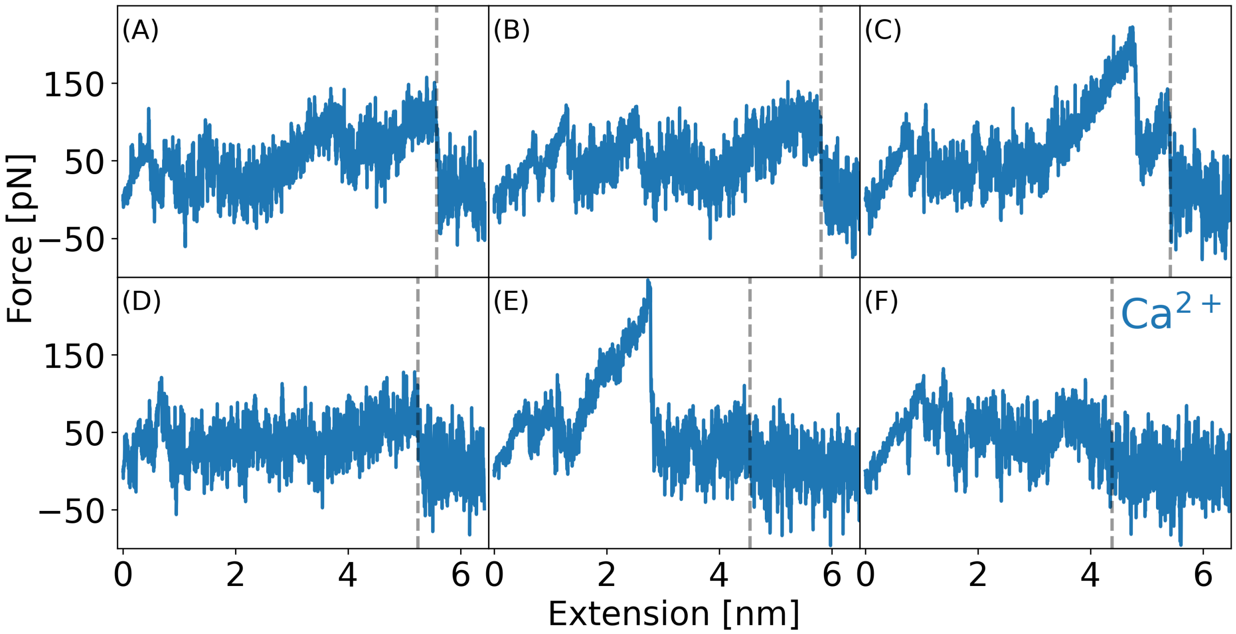


Figure S12: Force-extension profiles from simulations for $Ca^{2+}$ at a low simulation pulling speed of 0.1 m/s. The dashed vertical lines show the point where the DNA fully detaches from the mica surface.

#### 1.9 Diffusion coefficient

The diffusion coefficient is obtained from the in-plane mean square displacement (MSD) of the center of mass of DNA. The MSD is related to the diffusion coefficient by:

$$MSD = 4Dt, \tag{5}$$

Wherein $D$ represents the diffusion coefficient and $t$ is the time.

### 2 AFM Details

#### 2.1 Control experiments

Successful functionalization of an AFM cantilever tip with silane-PEG-mal (5 kDa) and subsequent poly-T ssDNA (30.5 kDa) is crucial for measurements on mica in different ionic solutions. As a control, AFM cantilever tips with PEG and DNA are first measured on mica in $H_2O$ (Figure S13) prior to experiments with DNA on mica in ion chloride solutions.

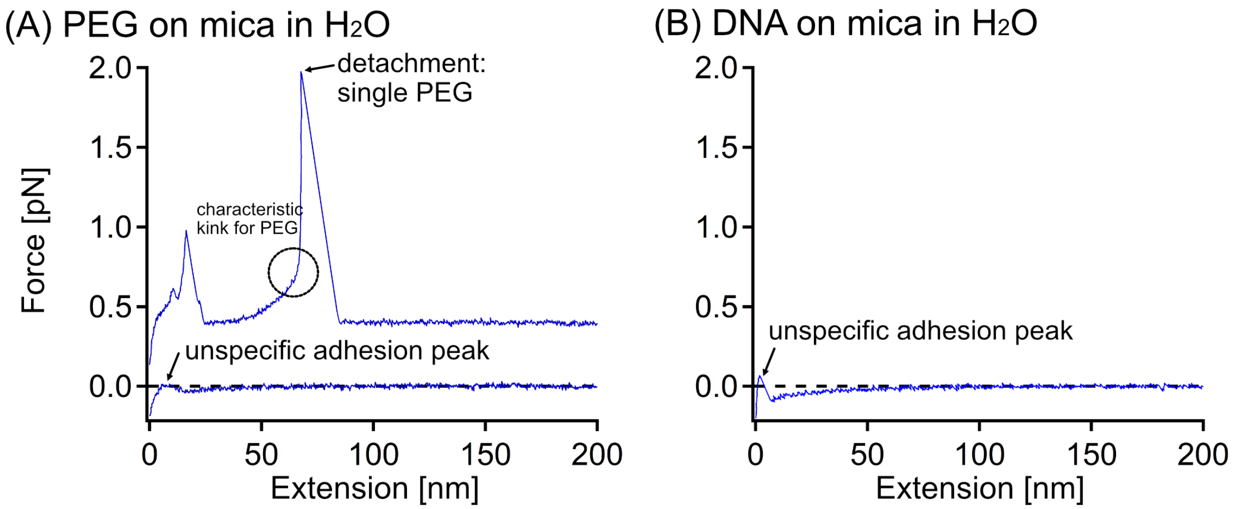


Figure S13: Representative force-extension curves of (A) PEG on mica in $H_2O$ (the force-extension curves are offset along the force axis in order to enhance their presentation), (B) DNA on mica in $H_2O$.

For PEG on mica in $H_2O$ either characteristic PEG stretching[4–7] events are observed or peaks at small extensions that account for unspecific adhesion between the AFM cantilever tip and the underlying surface (Figure S13A). Neither plateaus of constant force nor stick-slip events are observed. DNA on mica in $H_2O$ (Figure S13B) shows only unspecific adhesion peaks at small extensions.

#### 2.2 Loading rate determination

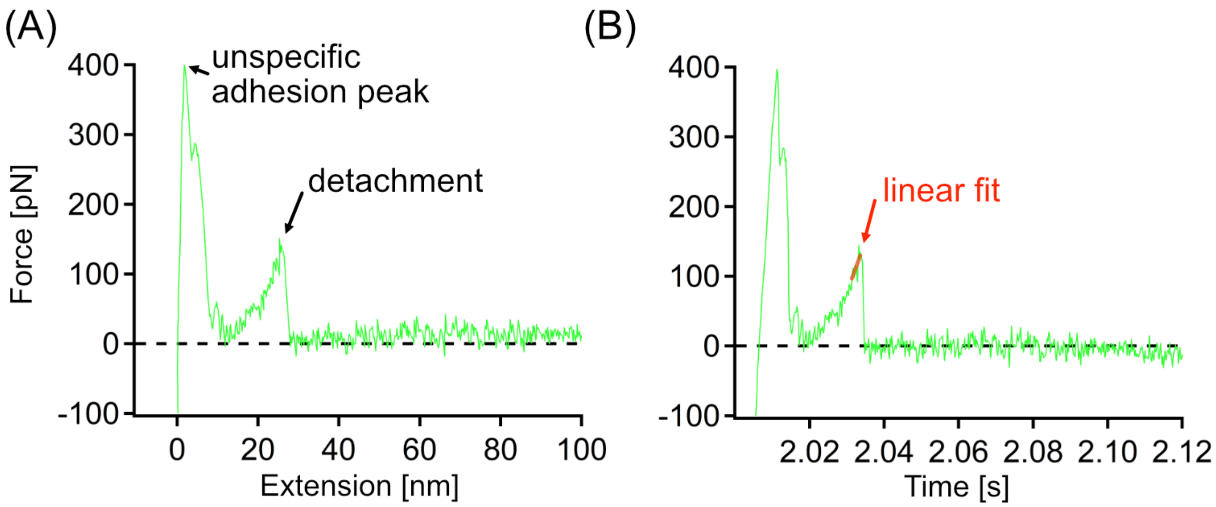


Figure S14: Determination of the loading rate based on force curves for DNA on mica in NaCl. (A) The last detachment event in stretching curves (represented by a force peak) is used to determine the detachment force of a single DNA molecule from mica. (B) Using the force-time curve the instantaneous loading rate is determined by fitting the last points of a stretching event prior to the last detachment event of the DNA from mica with a linear function, wherein the slope of the linear function corresponds to the loading rate.

#### 2.3 Overview of the experimental force-extension curves

For the analysis of the detachment of DNA from mica in different ion chloride solutions (LiCl, NaCl, KCl, CsCl), the force-extension curves are classified into two different categories of detachment events: stretching, including multiple stretching events, and plateaus according to Schwierz et al.[8] (Table S1 and Figures S15 and S16). Multiple plateaus are also observed, wherein each plateau possibly represents the detachment of a different DNA strand, as well as combinations of stretching and plateau events. In those cases the last event representing the detachment of the last DNA segment is evaluated for obtaining the detachment force.

Table S1: Overview of the number of curves, frequency and detachment forces for DNA on mica in different ion chloride solutions. In each force-extension curve the last detachment event has been taken to categorize the type of event into plateau or stretching events and to determine the respective detachment forces (see Figures S15 and S16). Finally, the total number of events and the respective detachment forces have been determined by merging the data for plateau and stretching events. The remaining curves do not show any single molecule events. They only show an unspecific adhesion peak, as given in Figure S13. For the forces values the mean values and the standard deviation of the respective distributions are given, respectively.

| Ion chloride solution (150 mM) | **LiCl** | **NaCl** | **KCl** | **CsCl** | **$MgCl_2$** | **$CaCl_2$** |
|---|---|---|---|---|---|---|
| **Total # curves** | **300** | **1100** | **300** | **100** | **900** | **1000** |
| # Stretching curves | 85 | 133 | 66 | 18 | 498 | 231 |
| Stretching curves [%] | 28 | 12 | 22 | 18 | 50 | 26 |
| Force (stretching) [pN] | 78 ± 55 | 84 ± 76 | 129 ± 151 | 148 ± 206 | 200 ± 129 | 178 ± 191 |
| # Plateau curves | 15 | 37 | 4 | 29 | 54 | 56 |
| Plateau curves [%] | 5 | 3 | 1 | 29 | 5 | 6 |
| Force (plateau) [pN] | 34 ± 12 | 64 ± 27 | 47 ± 43 | 30 ± 12 | 91 ± 38 | 82 ± 19 |
| **# Stretching & plateau curves** | **100** | **170** | **70** | **47** | **552** | **287** |
| **Stretching & plateau curves [%]** | **33** | **15** | **23** | **47** | **55** | **32** |
| **Force (stretching & plateau) [pN]** | **72 ± 53** | **79 ± 61** | **124 ± 149** | **75 ± 138** | **189 ± 127** | **159 ± 176** |

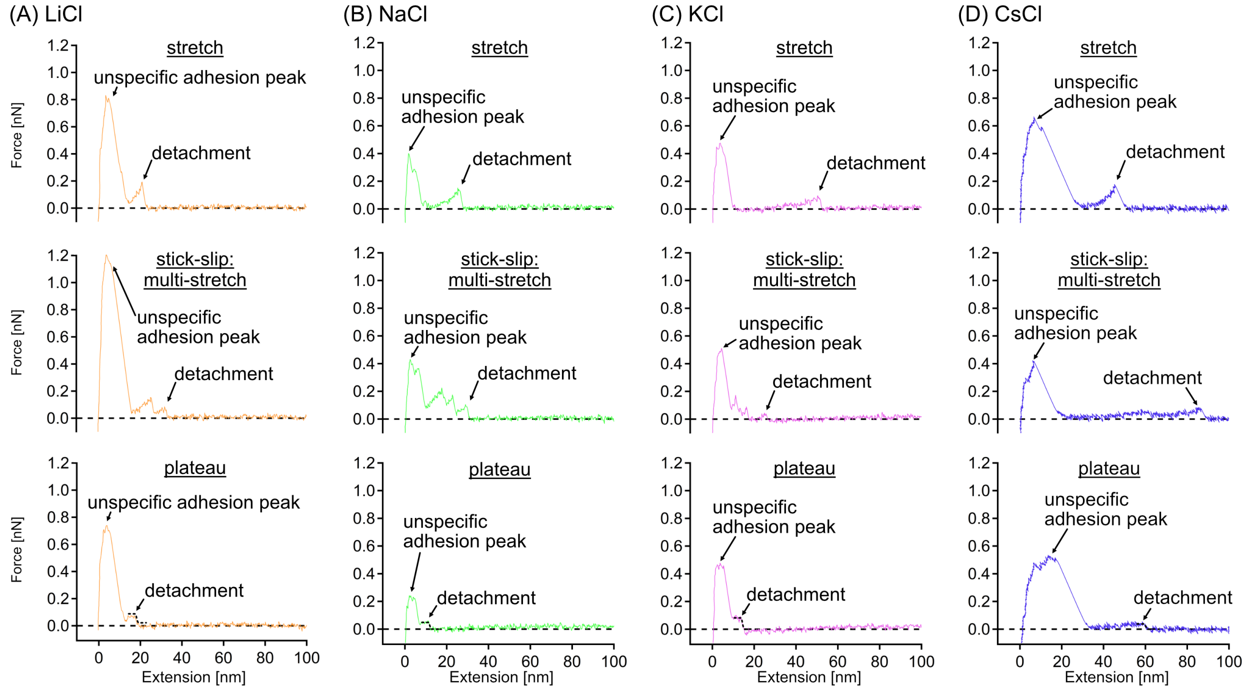


Figure S15: Examples of force-extension curves for DNA on mica in monovalent ion chloride solutions with different types of curves: stretches, multiple stretches and plateaus for (A) LiCl , (B) NaCl , (C) KCl and (D) CsCl. Final detachment events from which the detachment forces are taken are indicated by arrows. Stretching events (peak force as detachment force) or plateaus (detachment force taken from a sigmoidal fit to the end of the plateau) are clearly distinct from unspecific adhesion peaks at very small extensions resulting from an interaction of the whole AFM cantilever tip and the underlying mica surface.

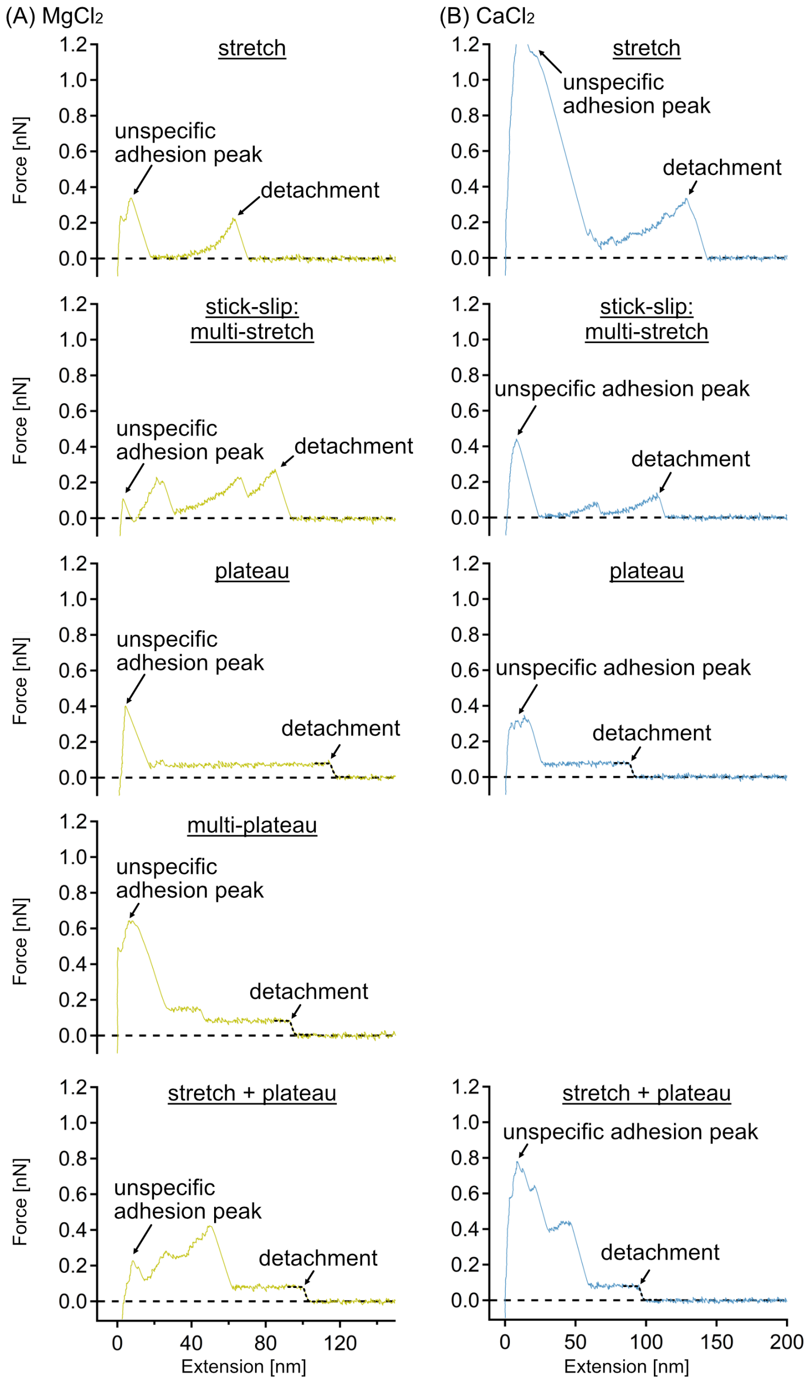


Figure S16: Examples of force-extension curves for DNA on mica in divalent ion chloride solutions with different types of curves: stretches, multiple stretches, plateau, multiple plateaus and combinations of stretches and plateaus for (A) $MgCl_2$ and (B) $CaCl_2$. Final detachment events from which the detachment forces are taken are indicated by arrows. These detachment events (peak force or plateau via a sigmoidal fit to the end of the plateau) are clearly distinct from unspecific adhesion peaks at very small extensions resulting from an interaction of the whole AFM cantilever tip and the underlying mica surface. $CaCl_2$ does not show any multi-plateau events.

For all types of cations, stretching is the most prominent type of single molecule event (Table S1). Stretching indicates a strong binding of the DNA to the mica surface, which then leads to a stretching of the polymer (PEG-DNA construct) by pulling.[9] The most common models to describe stretching are the freely jointed chain (FJC) or the wormlike chain (WLC) model.[10] PEG stretching in $H_2O$ is described by the two-state quantum mechanical freely rotating chain (TSQM-FRC) model.[7] Plateaus of constant force occur less frequently than stretching (Table S1). In the plateau region, DNA is in a stationary non-equilibrium with the mica surface during the desorption process.[11,12]

### 3 Results

#### 3.1 Average detachment forces from experiments and simulations

Table S2: Average detachment forces $\bar{\mathrm{F}}$ from simulations (v=10 $\mathrm{ms}^{-1}$) and experiments (v=1 $\mu\mathrm{ms}^{-1}$). Errors of the forces correspond to the standard deviation of the respective distribution.

| Ion type | $\bar{\mathrm{F}}_{\mathrm{sim}}$ [ pN ] | $\bar{\mathrm{F}}_{\mathrm{exp}}$ [ pN ] |
|---|---|---|
| $Li^+$ | $575 \pm 173$ | $72 \pm 53$ |
| $Na^+$ | $421 \pm 59$ | $79 \pm 61$ |
| $K^+$ | $404 \pm 69$ | $124 \pm 149$ |
| $Cs^+$ | $318 \pm 55$ | $75 \pm 138$ |
| $Mg^{2+}$ | $337 \pm 131$ | $189 \pm 127$ |
| $Ca^{2+}$ | $627 \pm 112$ | $159 \pm 176$ |

#### 3.2 Force distribution for $Mg^{2+}$ with and without high temperature pre-equilibration

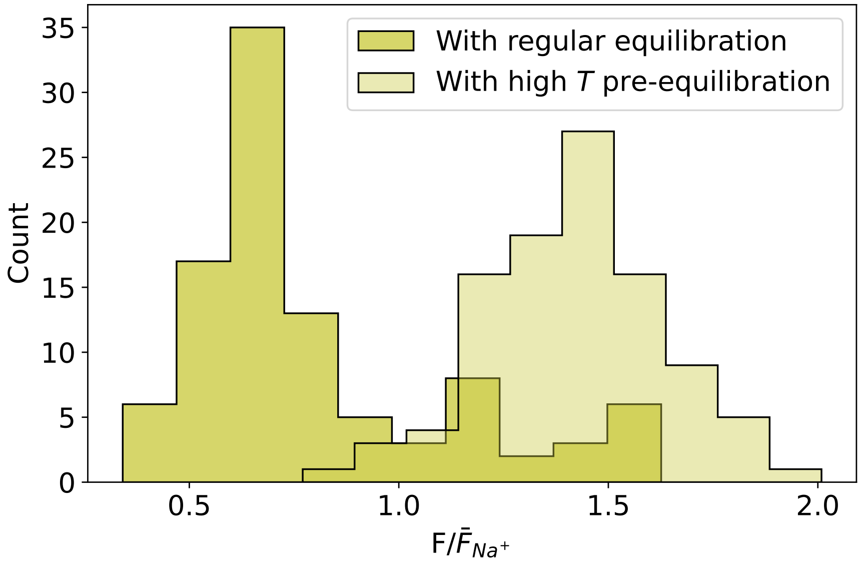


Figure S17: Comparison of force distribution for $Mg^{2+}$ with pulling after regular equilibration and with high temperature pre-equilibration. In the high temperature pre-equilibration the system was equilibrated first at 600 K for 200 ns. Subsequently temperature was reduced to 300 K and the regular protocol was followed as described in the main text. The regular equilibration result is the same as shown in Figure 4 K.
